# Dcifer: an IBD-based method to calculate genetic distance between polyclonal infections

**DOI:** 10.1101/2022.04.14.488406

**Authors:** Inna Gerlovina, Boris Gerlovin, Isabel Rodríguez-Barraquer, Bryan Greenhouse

**Affiliations:** EPPIcenter research program, Division of HIV, ID and Global Medicine, Department of Medicine, University of California, San Francisco, CA, USA

**Keywords:** genetic relatedness, identity by descent, genetic distance, ancestry, *Plasmodium*, polyclonal infection, microhaplotype

## Abstract

An essential step toward reconstructing pathogen transmission and answering epidemiologically relevant questions from genomic data is obtaining pairwise genetic distance between infections. For recombining organisms such as malaria parasites, relatedness measures quantifying recent shared ancestry would provide a meaningful distance, suggesting methods based on identity by descent (IBD). While the concept of relatedness and consequently an IBD approach is fairly straightforward for individual parasites, the distance between polyclonal infections, which are prevalent in malaria, presents specific challenges and awaits a general solution that could be applied to infections of any clonality and accommodate multiallelic (e.g. microsatellite or microhaplotype) and biallelic (SNP) data. Filling this methodological gap, we present Dcifer (Distance for complex infections: fast estimation of relatedness), a method for calculating genetic distance between polyclonal infections, which is designed for unphased data, explicitly accounts for population allele frequencies and complexity of infection, and provides reliable inference. Dcifer’s IBD-based framework allows us to define model parameters that represent interhost relatedness and to propose corresponding estimators with attractive statistical properties. By using combinatorics to account for unobserved phased haplotypes, Dcifer is able to quickly process large datasets and estimate pairwise relatedness along with measures of uncertainty. We show that Dcifer delivers accurate and interpretable results and detects related infections with statistical power that is 2-4 times greater than that of approaches based on identity by state. Applications to real data indicate that relatedness structure aligns with geographic locations. Dcifer is implemented in a comprehensive publicly available software package.

## 1 Introduction

Monitoring, effective control, and ultimately elimination of malaria can be accelerated by understanding the dynamics of malaria transmission including evaluation of interventions, identification of sources and sinks, and determining the drivers of sustained transmission. Given the substantial genetic diversity of malaria parasites, genomic data have the potential to illuminate important aspects of epidemiology (World Health Organization [2019]). Compared to viruses where mutations are the main source of variation and can be used directly to make temporal inferences, reconstructing transmission for recombining organisms with lower mutation rates requires a different approach. Since genetic recombination between malaria parasites occurs in the mosquito during person to person transmission, genetic relatedness can provide information on their shared ancestry and therefore transmission epidemiology at relevant timescales. Consequently, pairwise genetic distance as a measure of relatedness between infections may be more useful and detailed for answering epidemiologic questions than metrics based on comparison between populations (Wesolowski et al. [2018], Taylor et al. [2017], Tessema et al. [2019], Chang et al. [2019]). By assessing how closely related individual infections are, pairwise distance can also provide answers to questions such as whether particular infections were more likely to have been acquired locally or imported.

Due to co-infection and super-infection, individuals in endemic areas are often infected with multiple genetically distinct clones simultaneously.These polyclonal infections are the rule rather than the exception for *Plasmodium falciparum* in many endemic areas, even in relatively low transmission settings of sub-Saharan Africa (Roh et al. [2019]); polyclonality may be even more common for *Plasmodium vivax* (Koepfli et al. [2011], White et al. [2018]. Assessing genetic relatedness between polyclonal infections is more complicated both conceptually and methodologically than doing so for individual parasites. Obtaining phased genotypes of individual parasites from polyclonal infections would present a potential solution, but outside of single-cell sequencing this currently requires the use of statistical methods which are computationally intensive and may have limited accuracy in the absence of informative reference genomes, particularly when more than two clones are present (Zhu et al. [2019]). Even with phased genotypes, a unified summary of relatedness might be useful as a distance measure, so that it could be compared across pairs of infections which may be either monoclonal or have a higher complexity of infection (COI). Incorporating multiallelic genetic data, i.e. diverse loci with more than 2 variants, can improve estimation of relatedness between monoclonal infections (Taylor et al. [2019]) and may offer an even greater improvement over biallelic loci for polyclonal infections (Tessema et al. [2020]). Fortunately, current technologies make it feasible to efficiently amplify and sequence multiple diverse regions of the *Plasmodium* genome, generating multiallelic data for this purpose (Lerch et al. [2017], LaVerriere et al. [2021], Tessema et al. [2020], Aydemir et al. [2018]).

Much of the epidemiologically useful information contained in relatedness measures lies in detecting shared ancestry; there is therefore interest in estimating the proportion of genomes which are identical due to descent. Currently available methods based on identity by descent (IBD) for *Plasmodia* are developed for monoclonal infections or are adapted from human genetics: hmmIBD is designed for monoclonal infections and can incorporate multiallelic as well as biallelic loci (Schaffner et al. [2018]); isoRelate is able to accommodate polyclonal infections (Henden et al. [2018]), but is limited to biallelic loci and has unclear applicability to infections with COI > 2 since it is based on the diploid model. With no existing IBD-based methods to infer a degree of shared ancestry from polyclonal infections using multiallelic data, various suboptimal workarounds are generally employed. For example, some studies have attempted to infer a “dominant strain” from polyclonal infections using within host allele frequencies, while others have excluded polyclonal infections from the analysis altogether. Depending on the proportion of infections which are polyclonal, such procedures may grossly underutilize data or introduce bias to the analysis due to informative missingness. Alternatively, a simple identity by state (IBS) approach has been used (Tessema et al. [2019], Atuh et al. [2021]); it is convenient and fast but has extensive drawbacks as it produces similarity measures that are not easy to interpret and address relatedness only indirectly (Taylor et al. [2019]).

To fill the methodological gap, we introduce Dcifer (distance for complex infections: fast estimation of relatedness), a method employing IBD to estimate the level of common ancestry between polyclonal samples. It allows for unphased multiallelic data such as microsatellites or microhaplotypes as well as SNPs, explicitly takes into account COI and population allele frequencies, and does not require densely spaced or linked markers. Focusing on interhost relatedness, we developed a working model that allowed us to define an estimator with desirable statistical properties and formal inference. As the method provides a probabilistic solution to the multitude of possible underlying phased genomes, we used a unified mixed radix incrementing combinatorial algorithm for its implementation as a comprehensive [R software package](https://github.com/innager/dcifer). Finally, we assessed the performance of Dcifer for estimating relatedness between *P. falciparum* infections using simulations and empirical data.

## 2 Methods

Consider two infections with COI of *n_x_, n_y_* and a panel of *T* multiallelic markers. At each locus *t, t* = 1, …, *T*, there is a set *A_t_* = {*a*_*t*,1_, …, *a_t,K_t__*} of possible alleles. For convenience, we can arbitrarily order the alleles and map them to the corresponding population allele frequencies 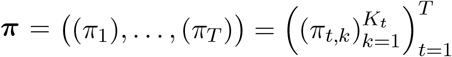. We assume that the underlying population allele frequencies are the same for both infections.

### 2.1 IBD model for two haplotypes

To build up a model for relatedness between polyclonal infections, we first consider two haplotypes. Let sequences of random variables *X* = (*X*_1_, …, *X_T_*), *Y* = (*Y*_1_, …, *Y_T_*) represent these haplotypes, and let (*IBD*_1_, …, *IBD_T_*) be a sequence of independent identically distributed random variables, where *IBD_t_* ~ *Bernoulli*(*r*) and parameter *r* describes the level of relatedness of the two haplotypes (Taylor et al. [2019]). Let *X_t_* ~ *P_t_*, where *P_t_* is a categorical distribution with values in *A_t_* and corresponding probabilites *π*_*t*,1_, … *,π_t,K_t__*, *X_t_* ⊥ *IBD_t_*; let *Y_t_* be a random variable such that

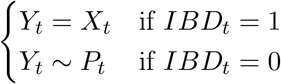

Note that *X_t_* and *Y_t_* are interchangeable in this setup, and the joint distribution of *X_t_* and *Y_t_* (marginal and conditional on *IBD_t_*) would not change if they were switched. While *IBD_t_* are i.i.d., (*X_t_, Y_t_*) are marginally independent but not identically distributed since *P_t_* is different for each *t*.

In this model, realizations of *X* and *Y* could be observed (e.g. if they represent monoclonal infections and there is no genotyping error) but *IBD*’s are unobservable. In contrast, 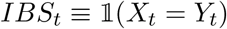 are directly observed if *X* and *Y* are observed.

### 2.2 Working model for polyclonal infections

Let an *n_x_* × *T* matrix ***X*** and *n_y_* × *T* matrix ***Y*** represent two polyclonal infections with COI of *n_x_* and *n_y_*, with rows of the matrices referring to haplotypes and columns to loci. Thus, *X_i_* = (*X*_*i*,1_, …, *X_i,T_*), is an *i*’th haplotype of the first infection, and a column *X*_1,*t*_, …, *X_n_x_,t_* is a sequence of random variables with values in *A_t_* representing alleles for all the haplotypes at a locus t. Let *S_x,t_* = {*X*_1,*t*_, …, *X_n_x_,t_*} denote a multiset (a collection of elements that are not necessarily distinct) of unordered elements of a *t*’th column of ***X*** and let *U_x,t_* = *Supp*(*S_x,t_*) = {*a_k_*: *a_k_* ∈ *S_x,t_*} be a set of unique elements in that column; *S_y,t_*, *U_y,t_* are defined similarly. For realizations of *S_x,t_* and *U_x,t_* we will use notation *s_x,t_* and *u_x,t_* (*s_x,t_* and *s_y,t_* are not observed, but *u_x,t_* and *u_y,t_* are). The model assumes no genotyping error, and the sequences of sets ***u**_x_* = (*u*_*x*,1_, …, *u_x,T_*) and ***u**_y_* = (*u*_*y*,1_, …, *u_y,T_*) are observed data, for which Dcifer is designed.

There are 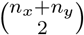 pairs of malaria strains that can be related. To differentiate *IBD_t_* and parameters of their distributions for different pairs, let *IBD_x_i_,x_j_,t_, IBD_y_i_,y_j_,t_*, and *IBD_x_i_,y_j_,t_* refer to a pair within first infection, a pair within second infection, and a between-host pair respectively, and, similarly, let *r_x_i_,x_j__*, *r_y_i_,y_j__*, and *r_x_i_,y_j__* denote corresponding relatedness parameters. If we are only interested in between-host relatedness (which may be the case for many practical applications), we might formulate the goal as “estimating interhost relatedness adjusted for intrahost relatedness”, which would condense 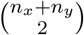 parameters into some lower-dimensional summary. Usefulness of adjusting for intrahost relatedness can be illustrated by considering a case where an extra haplotype *X*_*n*_*x*_+1_, very closely related to an existing one (say, *X*_1_, with *r*_*x*_1_,*x*_*n_x_*+1__ = 0.99), is added to one of the infections. That would result in essentially doubling *X*_1_’s contribution 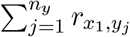 to the sum 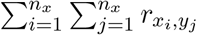 of all interhost relatedness parameters, as well as increasing COI (recall that *n_x_* is defined as a number of distinct haplotypes). With such goal as our scientific question, we introduce a simplifying assumption of no intrahost relatedness, which projects a realistic model of unconstrained intrahost and interhost relatedness parameters onto a much smaller model space and allows us to make the problem tractable while aiming to arrive at the same summary estimate as we would if we were able to estimate all the parameters in a bigger model.

For each pair of strains in two infections, e.g. *i*’th strain in the first sample and *j*’th in the second, let *X_i,t_, Y_j,t_*, and *IBD_x_i_,y_j_,t_* be the random variables as defined in Section 2.1. Then, for the working model for polyclonal infections, we introduce the following assumptions:

1. *r_x_i_,x_j__* = 0 for all *i, j* = 1, …, *n_x_*, *i* ≠ *j*, *r_y_i_,y_j__* = 0 for all *i, j* = 1, …, *n_y_*, *i* ≠ *j* (no intrahost relatedness);
2. *IBD_x_i_,y_j_,t_* ⊥ *IBD_x_k_,y_l_,t_* if *i* ≠ *k* or *j* ≠ *l* for all *t* = 1, …, *T* (all interhost *IBD* variables are independent at a given locus).

An important implication of these two assumptions is that any strain in one sample can be related to at most one strain in another: 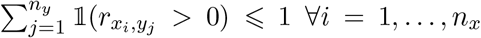 and 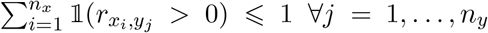. This can be proven by contradiction: since 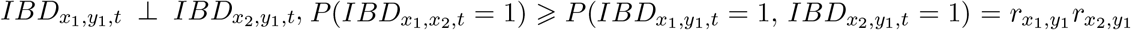. If *r*_*x*_1_,*y*_1__ > 0 and *r*_*x*_2_,*y*_1__ > 0, then *r*_*x*_1_,*x*_2__ > 0, which contradicts assumption 1. For further discussion on the model assumptions, see section S.1.

Since we can order strains within an infection arbitrarily, and in light of the constraints of the model, we order the haplotypes in two infections in such a way that *X*_1_ can only be related to *Y*_1_, *X*_2_ to *Y*_2_ and so on (Figure 1). In addition, we introduce *M* - the number of strain pairs that can be related, *M* = 1, …, min(*n_x_,n_y_*). Then, for brevity, we suppress some of the subscripts and use *r*_1_, …, *r_M_* for *r*_*x*_1_,*y*_1__, …, *r_x_M_,y_M__* and *IBD*_1,*t*_, …, *IBD_M,t_* for *IBD*_*x*_1_,*y*_1_,*t*_, … *IBD_x_M_,y_M_,t_* (note that parameters for all the other *IBD* variables are zero). The goal of Dcifer is to estimate parameters of the joint distribution of *IBD*_1,*t*_, …, *IBD_M,t_*. Let ***r*** = (*r*_1_, …, *r_M_*) denote an estimand, and 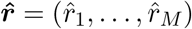 - its maximum likelihood estimator (MLE):

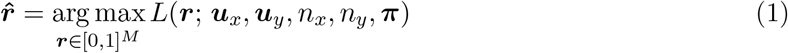

**Figure 1:**
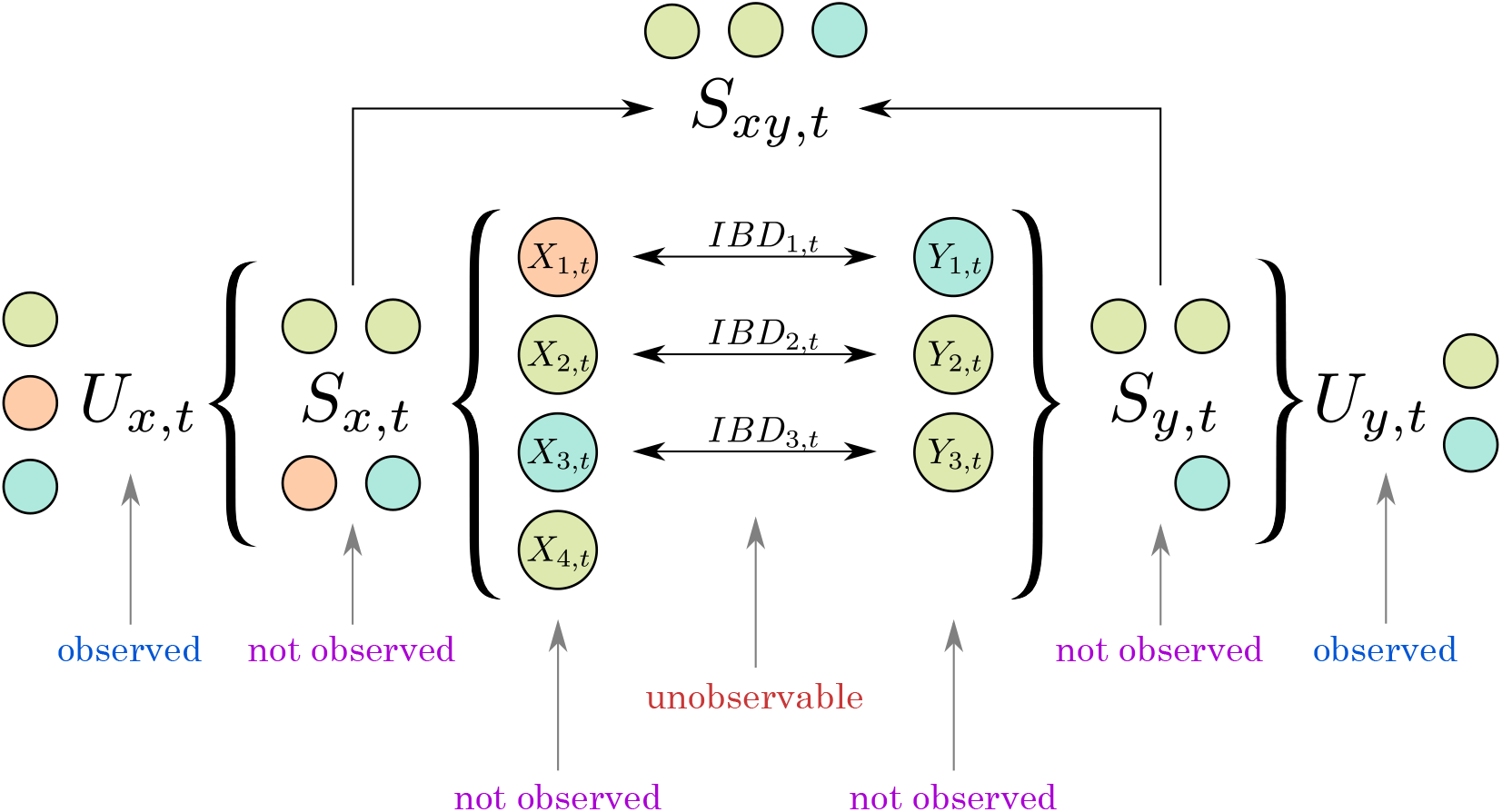
Working model presented at a single locus *t*: an example featuring *n_x_* = 4, *n_y_* = 3, and *M* = 3. Colors of the circles represent alleles; two clones in each infection have the same allele. *S_xy,t_* = *S_x,t_* ⋂ *S_y,t_* is a multiset of shared (non-unique) alleles at a locus *t*.

At each locus *t*, the likelihood *L*(***r**; **u**_x_, **u**_y_, n_x_, n_y_, **π***) needs to account for all the possible combinations of non-unique alleles in both samples (multiple haplotypes will have the same allele if COI is greater than the number of unique alleles). For one sample, this is done by considering a set of all multisets with given support and cardinality (all the *S_x,t_* that could have produced *U_x,t_*, see Figure 1); we denote a set of all multisets *s_x,t_* such that *Supp*(*s_x,t_*) = *u_x,t_* and |*s_x,t_*| = *n_x_* by *Q_x,t_*. *P*(*S_x,t_* = *s_x,t_*) can be calculated using a probability mass function of a multinomial distribution: the number of permutations of *s_x,t_* is equal to a multinomial coefficient (“assigning” alleles in *s_x_* to strains, or going from *S_x,t_* to (*X*_1,*t*_, …, *X_n_x_,t_*)), and allele frequencies correspond to event probabilities. Multiplicities of the multiset’s elements *a_t,k_* ∈ *A_t_*, or the numbers of strains having the same allele, are multinomial random variables. Adopting a short notation for this key component of the likelihood, let *g*(*s*^(*n*)^; *n*, (*π*)) denote a probability mass function for a multinomial distribution *Multinom*(*n, π*_1_, …, *π_K_*), where *s*^(*n*)^ is a multiset of cardinality *n*(|*s*^(*n*)^| = *n*) with elements from *K* categories, and (*π*) = (*π*_1_, …, *π_K_*) are probabilities for these categories; set 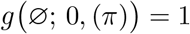. Next, for given *s_x,t_* and *s_y,t_* we divide their elements into three groups: shared alleles that are identical by descent (say *s*^(*m*)^), remaining alleles in *s_x,t_* (*s_x,t_*\*s*^(*m*)^), and remaining alleles in *s_y,t_* (*s_y,t_*\*s*^(*m*)^). The probability of each of these multisets is similarly calculated using multinomial distributions. Supplementary Section S.2 provides more details and builds up the likelihood from *M* =1 and *M* = 2. For a general case,

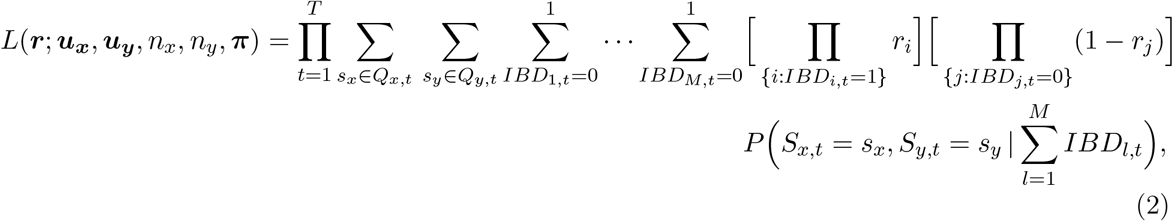

where

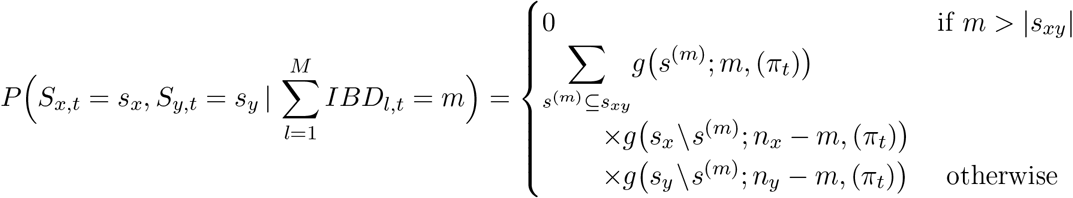

and *s_xy_* = *s_x_* ⋂ *s_y_*.

When *r*_1_ = *r*_2_ = … = *r_M_* = *r*, the likelihood reduces to

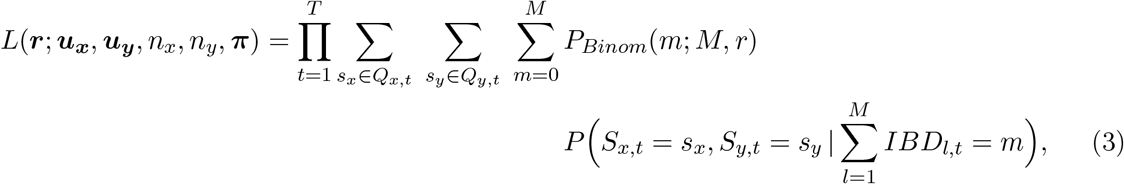

where 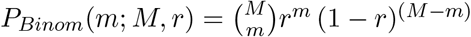.

While *IBD*_1,*t*_, …, *IBD_M,t_* are independent, 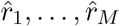 are not. This dependence stems from the fact that we do not observe ordered alleles at each locus (or, in other words, phased haplotypes). That also provides intuition for why 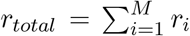 is estimated more accurately than individual *r_i_*’s: estimating 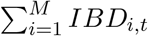 at a locus *t* is easier than estimating an actual binary sequence (*IBD*_1,*t*_, …, *IBD_M,t_*). Another useful observation is that the order of parameter values in ***r*** does not affect the value of *L*(***r**; **u**_x_, **u**_y_, n_x_,n_y_*, **π**), which can be taken into account when the likelihood is evaluated over a grid of ***r*** ∈ [0,1]^*M*^.

### 2.3 Implementation

Calculating the likelihood in (2) requires solving a number of combinatorial problems: finding all the collections of non-unique alleles at a locus that are concordant with observed alleles and COI, finding all the multisets included in a given multiset of shared non-unique alleles, and finding all the possible binary sequences with given constraints for *IBD* variables. These problems are solved with a unified mixed radix incrementing algorithm (Gerlovina [2022]), which is an extension of an algorithm to generate all *n*-tuples in Knuth [2011]. As the calculation traverses the combinations described above, multiple ***r*** = (*r*_1_, …, *r_M_*) sequences can be processed at each step, and thus the likelihood for a range of parameter values can be calculated in a single pass. With bounded parameter space, this allows for an efficient way to find MLE by simply calculating the likelihood for an *M*-dimensional grid of a desirable coarseness. The resulting support curve or surface can also be useful for inference, especially for procedures based on a likelihood ratio approach, such as testing various hypotheses or determining confidence regions. For a special case of *M* = 1, the log-likelihood can be calculated using Equation 5, which also admits fast calculation of the score and consequently numerical methods of solving the likelihood equation.

### 2.4 Inference

Along with an estimate of ***r***, Dcifer provides a support curve/surface - log-likelihood values for a grid of ***r*** values, which can provide a basis for various inferential procedures (for some intuition on the shape of that support curve, the effect of COI and population allele frequencies on it, and implications for the inference, see Supplementary section S.3). In our model, sample size is *T*, but different loci do not provide the same amount of information (recall that (*X_i,t_,Y_j,t_*), *t* = 1, …,*T* are independent but not identically distributed); their contribution can be associated with different measures, e.g. heterozygosity. Given these measures and the complexity of the estimator, methods relying on asymptotic approximations should be approached cautiously; still, as the sample size increases, precision of estimation increases as well.

For hypothesis testing and confidence intervals/regions, we consider common inferential approaches as applied to Dcifer: asymptotic normality, likelihood-ratio statistics, and resampling methods. There are common challenges that affect all three approaches: bounded parameter space [0, 1]^*M*^ with edge cases not only included but conceptually important, such as a null hypothesis *H*_0_: *r*_1_ = … = *r_M_* = 0 of infections being unrelated; for other cases, sampling distributions for different (even neighboring) parameter values on the interior of the support could be quite different for panels with even fairly large *T*. Still, some approaches might be better suited for Dcifer, and some may be chosen on the basis of convenience and computational efficiency. For Wald-type confidence intervals, observed Fisher information can be easily calculated numerically (and arguably preferred to expected Fisher information - Efron and Hinkley [1978]); likelihood-ratio-based CI, while asymptotically equivalent to Wald’s (Wald [1943]), are more robust as they are invariant to parameter transformation that could be used to make the support curve approximately quadratic at MLE (Meeker and Escobar [1995], Beale [1960], Cox and Hinkley [1979], pp. 342-343, Vander Wiel and Meeker [1990], Cook and Weisberg [1990]). Resampling methods include bootstrap and generating a null distribution for hypothesis testing. While there are many advantages this approach provides for finite sampling distributions not yet approaching normality, there is a caveat: if a centered sampling distribution at MLE is not close enough to that at the true value, the inference will be problematic. In addition, inverting quantiles of a bootstrap distribution for CI endpoints can lead to violating the bounds of the parameter space (see Supplementary section S.4), as can Wald CI. Simulated null distributions do not suffer from this problem but still rely on various assumptions and might be sensitive to misspecifications as demonstrated in the Results section. In contrast, likelihood-ratio confidence regions respect parameter bounds and do not require any additional model assumptions for hypothesis testing.

Likelihood-ratio-based inference is based on Wilks’ theorem (Wilks [1938]) and uses the likelihood ratio test statistic, which in the context of Dcifer hypothesis testing with *H*_0_ : ***r*** = ***r***_0_ can be written as 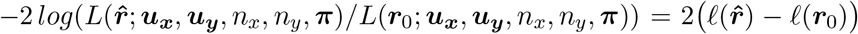, where ℓ(***r***) = *log*(*L*(***r***; ·)), and is approximated by chi-squared distribution *χ*^2^(*M*) with *M* degrees of freedom. The approximate 1 – *α* confidence region consists of the values

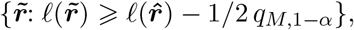

where *q*_*M*,1–*α*_ is a (1 – *α*)’th quantile of *χ*^2^(*M*). As the Wilk’s theorem does not apply to border cases, we specifically address these important cases (for which likelihood-ratio test is still most powerful Neyman and Pearson [1933]) and compare the corresponding distribution of the likelihood ratio statistic with *χ*^2^(*M*) that no longer approximates it. First, the test at the boundaries is one-sided while the chi-squared distribution implies two-sided tests. Accounting for that would mean dividing a p-value obtained from the chi-squared distribution by 2 or finding a corresponding critical value for the significance level *α*. Second, it turns out that even with this adjustment, the resulting p-value is still somewhat conservative, and, as shown in the Results section, the method has excellent error rate control.

### 2.5 Estimating the number of related strain pairs and *r_total_*

Parameters of the working model include *n_x_, n_y_*, and ***π***. *M*, the length of ***r***, can be considered a nuisance parameter. In addition, let 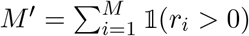 be a number of positively related strain pairs; unlike *M*, *M*′ can be a quantity of interest. Estimator 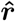 in Equation 1 assumes that the model is constrained by given values of *n_x_, n_y_*, ***π***, and *M*; the likelihood is calculated using these values. However, while *n_x_, n_y_*, and ***π*** are “external” to ***r*** and are provided or obtained through other processes, *M* is inherent to relatedness between two infections. Thus here we consider a less constrained model where *M* is not given. In this case, a trivial solution to estimating ***r*** would be to set *M* = min(*n_x_, n_y_*) since ***r***’s associated with different *M*′ ⩽ *M* ⩽ min(*n_x_,n_y_*) will only differ in the number of zeros (*r_i_* = 0). If we want to estimate *M*′ or 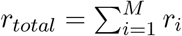, which are functions of ***r***, they can be similarly obtained from 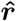 as 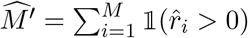 and 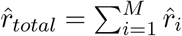.

In practical applications this trivial solution can incur high computational cost for higher min(*n_x_*, *n_y_*), and therefore we propose alternative estimators 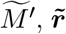, and 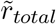 that use an iterative procedure with underlying calculation of 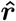 at each step. The first step is to set *M* = 1 and calculate 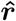, then at each consecutive step increment *M* and recalculate 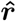 until it contains one zero 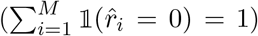 or until *M* = min(*n_x_,n_y_*). Accept 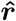 obtained at the final step as 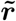, with 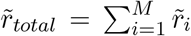 and 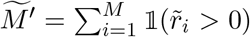. If *r_total_*, rather than (*r*_1_, …, *r_M_*), is of main interest, the computation time can be cut even further by assuming *r*_1_ = … = *r_M_* = *r* and using Equation 3 to calculate the likelihood. In this case we propose yet another set of estimators 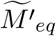 and 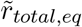, where 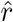 is calculated for all *M* = 1, …, min(*n_x_,n_y_*), and 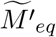 is the value of *M* that produced the highest maximum likelihood. Then 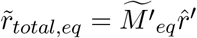, where 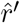 is an MLE at 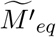.

### 2.6 Simulations

Our simulations are based on previously published single nucleotide polymorphism (SNP) and microhaplotype panels (Daniels et al. [2008], Jacob et al. [2021], Tessema et al. [2020]). To include genotyping errors in simulations, we devised a “miss-and-split” model with parameters *ϵ* and λ:

1. False negatives: one of *k* present alleles (drawn with probabilities 1/*k*) has zero probability of being missed; the remaining *k* – 1 alleles can be missed with probability *ϵ*. Let *K* be a number of alleles remaining present after this step; then *E*(*K*) = 1 + (*k* – 1)(1 – *ϵ*).
2. False positives: draw a number *N_add_* ~ *Pois*(λ) of added alleles (“splitting” event) for each non-missing allele; subsequently draw *N_add_* alleles from *K* – 1 alleles with replacement. In the final “observed” data, an allele is considered present if selected by at least one of the splitting events.

Note that *P*(*N_add_* ⩾ 2) is very small for reasonably small λ’s.

For analysis procedures that involve estimating COI and population allele frequencies prior to Dcifer, we use naïve COI estimation with an offset *c* (COI determined by a locus with *c*’th greatest number of detected alleles), where *c* depends on the number of loci, for multiallelic panels, and THE REAL McCOIL method for biallelic SNP panels (Chang et al. [2017]). For allele frequencies, we use COI-adjusted estimation (see Supplementary Section S.5), which is important for polyclonal infections. Failure to adjust for COI can lead to overestimating heterozygosity and, consequently, relatedness parameters.

## 3 Results

The main goal of Dcifer is to estimate parameters describing relatedness between infections, and this estimation requires values of the other parameters in the model. These external parameters represent COI (*n_x_*, *n_y_*) and population allele frequencies (***π***), which can be known (e.g. in simulations), estimated from data, or otherwise specified. Dcifer is implemented in a software package that takes raw data on the alleles detected at each locus (biallelic or multiallelic) in *Plasmodium* infections, allowing for missing data, along with COI and population allele frequencies. In simulations, we assess the performance of Dcifer when data have no genotyping error and COI and ***π*** are known, as well as in the presence of genotyping error with COI and ***π*** estimated from these data. We also evaluate how sensitive Dcifer is to misspecification of these external quantities and to assumption violations. We start with a case when only one pair of strains can be related (*M* = 1), since it can be used to quickly identify related infections in a large dataset, and later proceed to the general case. Finally, we apply Dcifer to analysis of real data, where COI and allele frequencies are estimated from the data.

### 3.1 Dcifer produces accurate and interpretable estimates of relatedness

Unlike IBS metrics that simply measure similarity between infections comparing detected alleles, Dcifer aims to produce more interpretable results by estimating parameters that represent IBD and thus separating shared ancestry and chance as underlying reasons for alleles matching between two infections. To evaluate the performance of this method in comparison to IBS approach (we used Jaccard similarity coefficient as an example), we simulated genetic data for infection pairs with different degrees of relatedness (induced on a single pair of strains between two infections) and COI, based on previously published single nucleotide polymorphism (SNP) and microhaplotype panels (Daniels et al. [2008], Jacob et al. [2021], Tessema et al. [2020]). Across various values of COI, Dcifer estimates were concentrated around the true values of the parameter while IBS results were not (Figure 2, shown for a panel of 91 microhaplotypes with no genotyping error and known COI and allele frequencies). As COI increased, Dcifer estimates became more variable but remained centered around the true values and maintained some degree of separation, whereas IBS results shifted and overlapped considerably more. For these simulations, separation between results for completely unrelated (*r* = 0) and related infections, quantified in ROC curves, indicated considerable gain in accuracy by Dcifer compared to the IBS metric across the range of COI, especially for lower degrees of relatedness (Supplementary Figure S.1). For example, Dcifer estimates of sibling-level relatedness (*r* = 0.5) remained readily distinguishable from those of unrelated infections even for fairly high COI.

**Figure 2:**
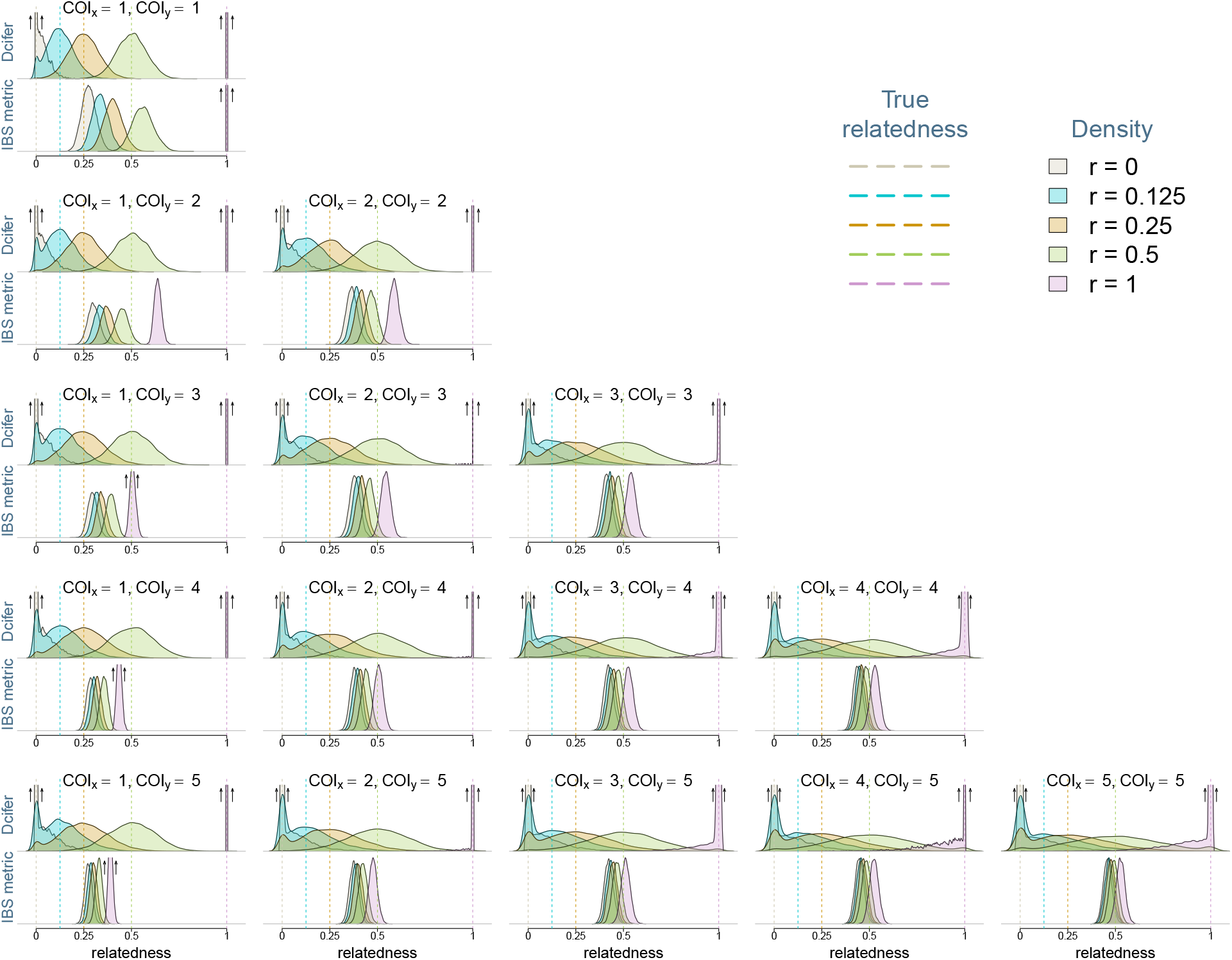
Densities of Dcifer relatedness estimator 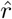 and IBS similarity metric results obtained from data simulated using a panel of 91 microhaplotypes (Tessema et al. [2020]). Simulations were performed for five values of *r* and for COI combinations ranging between 1 and 5; true values of COI and population allele frequencies were used for Dcifer. Upward arrows indicate highly concentrated distributions with density values extending above the plot range (note: y-axis scales are different for the two methods).

When the task of detecting related infections is approached in practice, there are additional issues to be considered because neither COI nor population allele frequencies are known, genetic data often contain genotyping errors, and the extent of these errors is unknown. Supplementary Figure S.2 illustrates how results changed when genotyping error was included in the simulations, and estimates of COI and population allele frequencies, and not their true values, were used as inputs to Dcifer. IBS results shifted to the left more or less uniformly; since distributions for different values of *r* were so tightly concentrated and close together, apparently small shifts were significant compared to the differences between these distributions. Dcifer estimates also shifted to the left, but, relative to differences in sampling distributions, the shifts were smaller than those for IBS. The fact that the shifts were more pronounced for larger values of *r* is explained by genotyping errors breaking up some of the relatedness between infections.

### 3.2 Greater power of hypothesis tests using Dcifer vs. IBS

One way of detecting related infections along with a measure of uncertainty (e.g. p-value) is to compare Dcifer relatedness estimates or IBS similarity results with their corresponding null distributions (*H*_0_: *r* = 0), which can in theory be obtained by simulating a large number of unrelated infections. To evaluate performances of Dcifer and the IBS metric, we calculated false positive rates (FPR) and power of tests with significance level *α* = 0.05 for different types of genetic data across a range of COI. Genetic data were simulted with genotyping errors; they were incorporated into simulated “null” distributions as well. Distributions for *r* = 0 are different for different COI and therefore a separate null distribution was generated for each COI pair combination; the effect of COI on such distributions was substantial for IBS (Supplementary Figure S.3). Relatedness estimates for each pair of infections were then compared to a rejection cut-off determined by a null distribution corresponding to their estimated COI, and FPR and statistical power were subsequently calculated. In addition to the complexity and computational costs associated with generating a null distribution, this approach relies on a number of assumptions such as COI, allele frequencies, and the error model and its parameters, which are all subject to misspecification.

As a welcome alternative, Dcifer offers another inferential approach based on the likelihood ratio, which does not require any additional information (i.e. does not require generating a null distribution) and has essentially no computational overhead. Figure 3 compares hypothesis testing results for IBS, using simulated reference distributions, and Dcifer, using likelihood-ratio p-values adjusted for one-sided tests. For both methods, FPR was mostly at or below the nominal significance level *α* across different simulations of COI and genotyping panels, with Dcifer close to *α*. Statistical power, however, varied considerably. As expected, higher values of relatedness were detected with greater power, increasing the number or diversity of loci increased power, and higher COI led to lower power. Across all simulations, Dcifer consistently demonstrated greater power to detect related infections than the IBS metric, with differences particularly notable for polyclonal infections. For example, with a 91 microhaplotype panel, the power to detect half-siblings (*r* = 0.25) in a pair of infections with COI of 2 was 0.81 for Dcifer and 0.43 for the IBS metric; with 455 microhaplotypes and COI of 5 that power was 0.88 and 0.22 respectively. While Supplementary Figures S.1 and S.2 would suggest that there is still some separation between distributions for different values of *r* for the IBS metric results, which would be expected to improve with increasing the number of loci, its performance was remarkably poor, having very low power for larger COI and *r* < 0.5 even with highly informative panels. This reflects the fact that for tightly concentrated distributions of IBS results, the difference between cut-offs associated with different assumed null distributions is critical, and consequently misspecification of COI or an error process had a deleterious effect on either FPR or power (Supplementary Figure S.3). The likelihood-ratio based approach performed very similarly to the one based on null distributions for Dcifer, evidencing this as a preferred approach for the reasons described above (Supplementary Figure S.4).

**Figure 3:**
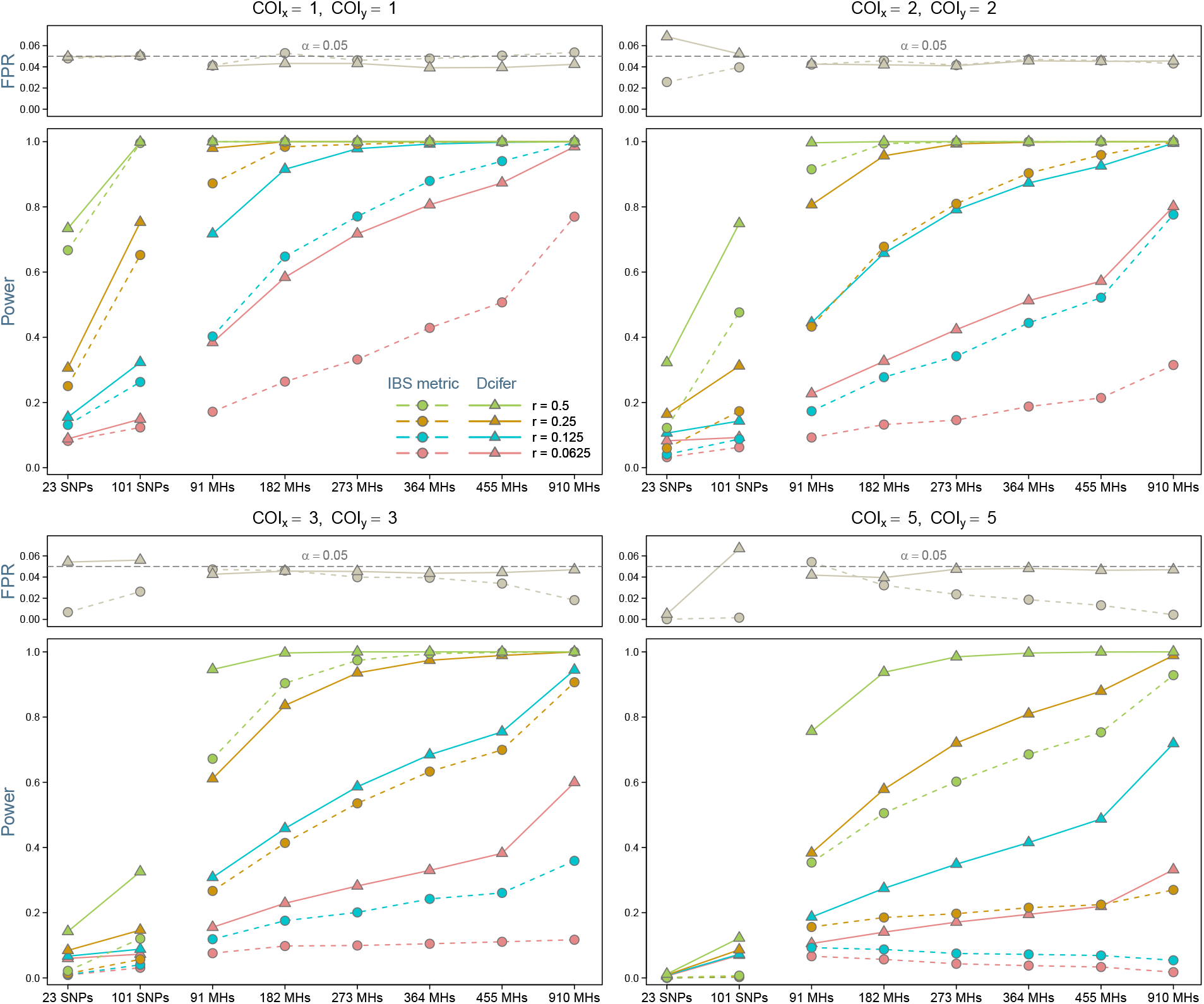
Detecting related infections. False positive rate and statistical power of a test *H*_0_: *r* = 0 at significance level *α* = 0.05 are shown. Simulations included genotyping error with fixed error model parameters; COI were estimated from these data. For simulated null distributions, we varied error model parameters since they would not be normally known. Single nucleotide polymorphism (SNP) and microhaplotype (MH) panels were used as a basis for simulations.

### 3.3 Dcifer provides likelihood-ratio-based confidence intervals

The Dcifer likelihood-ratio-based approach allows for calculating *M*-dimensional confidence regions (where *M* is the number of related pairs) - or, in a case when only one pair of strains is assumed to be related between two infections, confidence intervals (CI). Figure 4 shows CI’s for a range of true *r* values and COI. Infections were simulated using microhaplotype panels with various number of loci. As expected, CI’s were narrower for panels with more loci. In general, the intervals were narrower near endpoints (*r* = 0 and *r* = 1) and wider in the midrange. Interestingly, the least COI in the pair (min(*n_x_,n_y_*)) had a greater effect on the CI than the sum of COI (*n_x_* + *n_y_*); this can be seen in more rapid widening of the intervals from left to right than from top to bottom of the figure. With large numbers of diverse loci, CI stayed narrow even for higher COI. Coverage for these CI was around 1 – *α* = 0.95, and consistently higher for endpoints, indicating that CI for these endpoints were conservative, even taking into account the one-sided nature of such intervals (Supplementary Figure S.5; also demonstrated by FPR in Figure 3).

**Figure 4:**
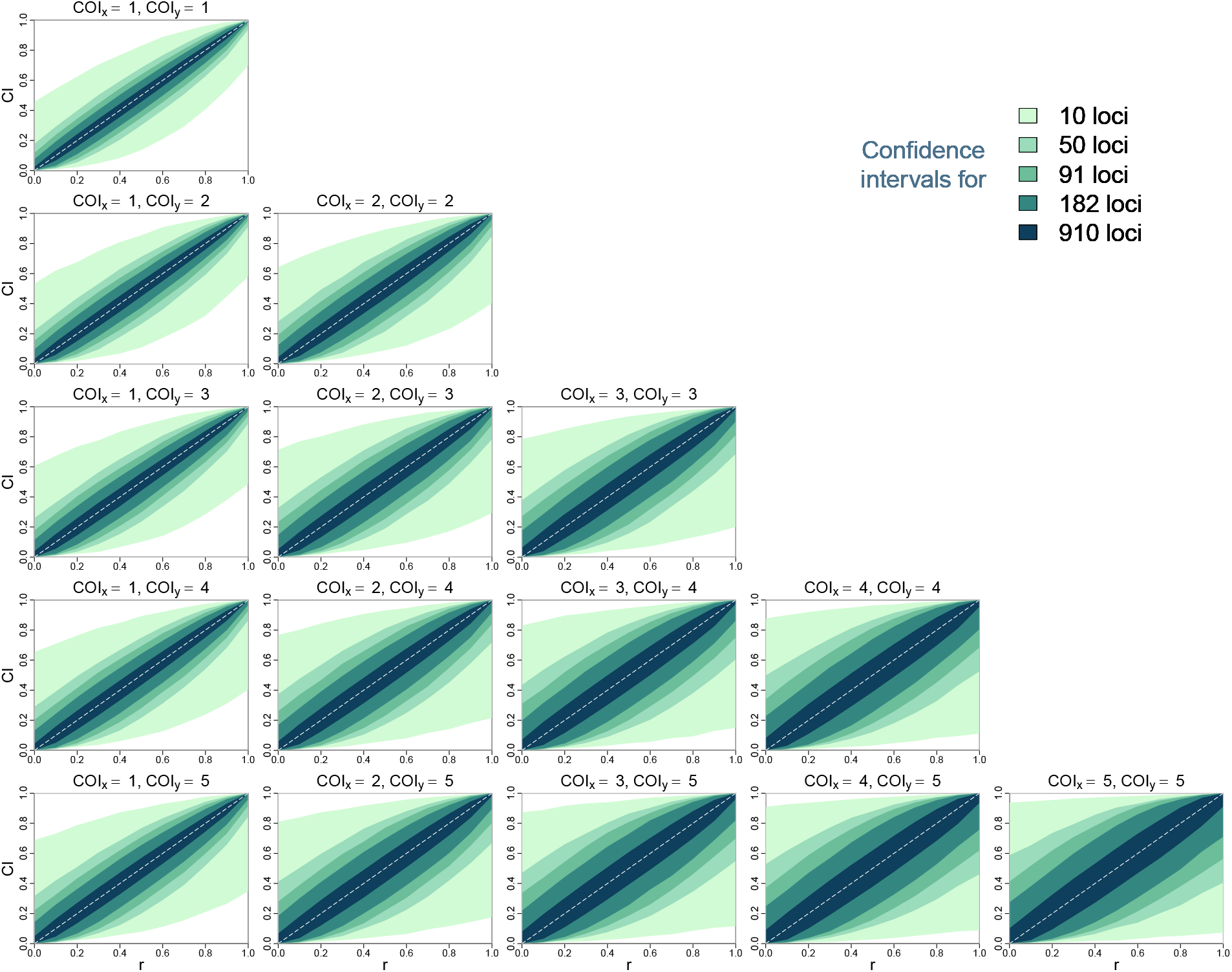
95% confidence intervals (CI) for relatedness estimates using the likelihood ratio produced by Dcifer.

### 3.4 Allowing for multiple pairs of strains to be related

So far we have only presented results for a single related pair of strains between two infections (*M* = 1) regardless of COI. When we allow that multiple pairs of strains may be related, Dcifer produces a corresponding number of estimates - one for each pair. To accurately estimate multiple relatedness parameters without any additional assumptions, a large number of diverse loci is needed; otherwise, there is a lot of variation in the individual estimates (see an example in Supplementary Figure S.6). However, while estimation of individual relatedness parameters was challenging, their sum *r_total_* was estimated more accurately even with a lower number of loci (which can be seen in the contour plots of Supplementary Figure S.6). Supplementary Figure S.7 shows likelihood surfaces for two-dimensional parameters (*M* = 2), where we can clearly see the ridge along 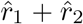; that value is close to the true sum even when the individual estimates are further away from (*r*_1_, *r*_2_).

If the goal is to estimate overall relatedness between two infections, we suggest *r_total_* as a more identifyable and useful quantity than (*r*_1_, …, *r_M_*). To estimate *r_total_* and the number of positively related strain pairs *M′*, we used the procedure described in Section 2.5 and compared the estimates obtained from two approaches: (1) with “equality assumption” *r*_1_ = … = *r_M_* (estimators 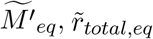) and (2) without it (estimators 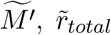). First, we validated the stopping rule for the second approach (when *r_i_* are not assumed to be equal), confirming that incrementing *M* past the iteration that estimates one of *r_i_*’s to be 0 only appended additional 0’s to MLE in most cases. Next, we compared the two approaches and assessed the accuracy of the corresponding estimators. For each simulated pair of infections, we first randomly generated *r*_1_, …, *r_M′_*, 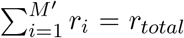 for given *r_total_* and *M′*. The estimates were compared across a grid of COI, *r_total_*, and *M′*. Figure 5 shows illustrative examples of these comparisons: in 5a, *M′* is changed while COI and *r_total_* are fixed, in 5b *r_total_* is changed, and in 5c COI is changed. Distributions of 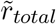 and 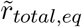 were quite similar, so the equality constraint had a very limited effect on the overall relatedness estimates. There was more difference between 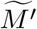 and 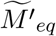, but, importantly, these differences did not significantly affect *r_total_* estimates. An effect of varying *M′* on the distributions of 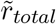 and 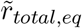 was small, while lower COI resulted in more accurate estimates. Higher *r_total_* made for more accurate estimation of *M′*, as it eliminated lower values incompatible with *r_total_* estimates. It is worth noting that simply increasing dimensionality of the grid of relatedness values to evaluate over can become unfeasible for larger *M*, so the grid would have to be coarsened to accommodate, which in turn would affect precision. No such limitation exists for the fast “equal *r_i_*” approach as it estimates a single parameter.

**Figure 5:**
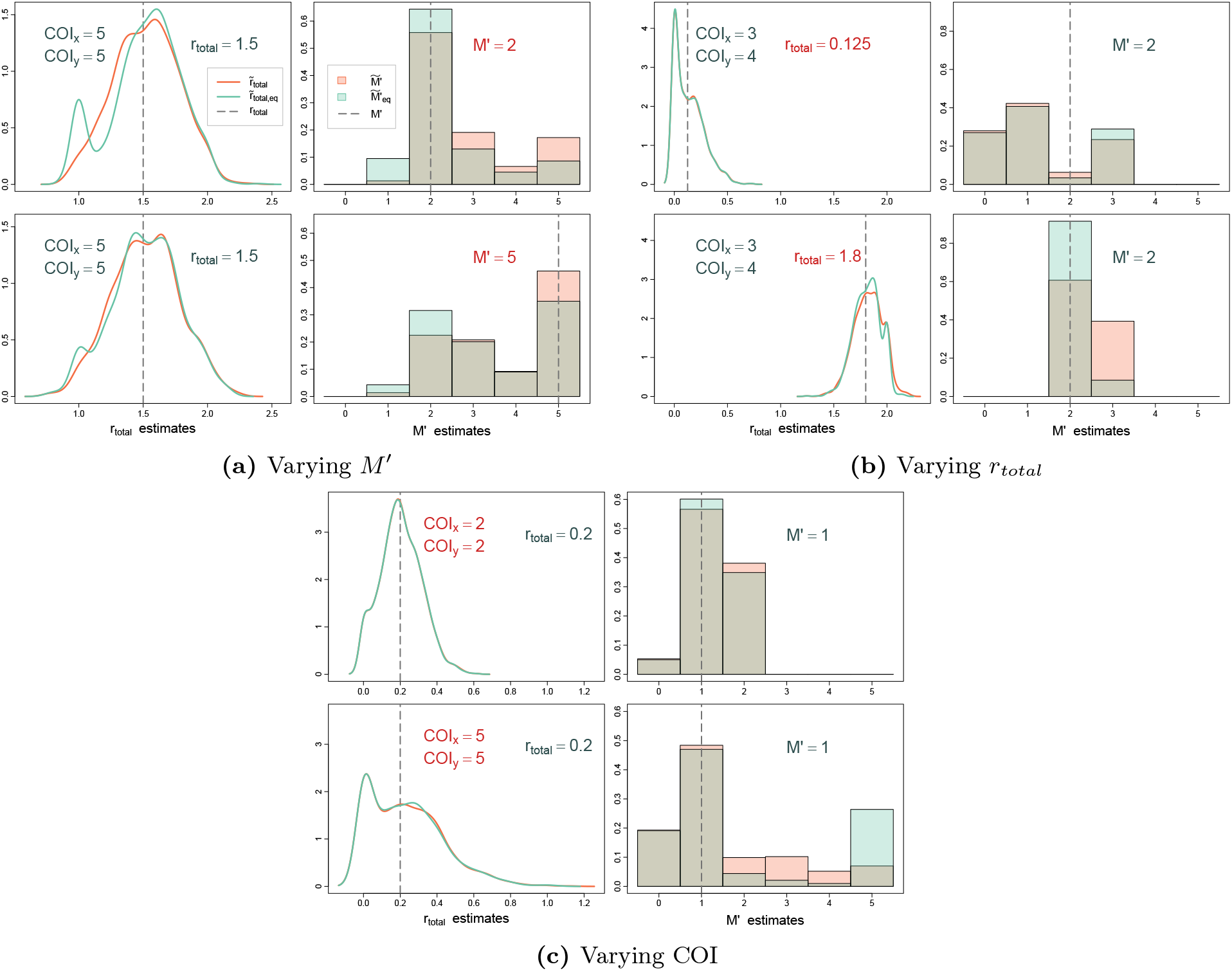
Estimation of *r_total_* and *M′* with and without equality assumption *r*_1_ = … = *r_M_*. Densities of 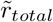 (no assumption) and 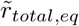 (with assumption) are shown on the left, and probabilities for the values of 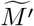 and 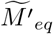 are shown on the right. Quantities highlighted in red indicate those varied between the top and bottom simulations within each panel. Simulations were performed using a panel of 91 microhaplotypes.

### 3.5 Misspecifications and assumption violations

In data analysis, COI and population allele frequencies are usually unknown and need to be estimated from data. Allele frequencies can often be estimated from sufficiently large datasets (e.g. over 100 samples) and as such, their estimates are often fairly stable; some implications of their misspecifications are discussed in Section S.3 of Supplementary materials. COI estimation, however, relies on a smaller amount of information, resulting in greater variability of the estimates and more frequent misspecifications. Fortunately, Dcifer appeared to be relatively robust to COI misspecifications, especially for less complex infections (Supplementary Figure S.8). Even for higher COI, relatedness estimates were fairly close to the true value in the neighborhood of the correct COI.

Next, we address our working model and its defining assumption of no intrahost relatedness. To assess how violating this assumption affects interhost relatedness estimation, we compared three simple scenarios: one with no intrahost relatedness and two where different strains within one of the samples are siblings (*r* = 0.5) - see the diagrams in Supplementary Figure S.9. Note that in scenario 2 (Supplementary Figure S.9b), there are two interhost pairs that are related (strains *X*_1_-*Y*_1_ and *X*_2_-*Y*_1_) but that “extra” relatedness is only a consequence of the induced *X*_1_-*X*_2_ sibship and does not add anything to our quantity of interest. All simulations included genotyping error, and processing involved estimating COI and population allele frequencies. Relatedness estimates for all three scenarios were very similar (Supplementary Figure S.10), confirming that in simple cases of intrahost relatedness, the working model estimates interhost relatedness without significant biases.

### 3.6 Applications to empirical data

We applied Dcifer to a small dataset that has 87 microhaplotypes and consists of samples obtained from patients presenting with malaria from two health facilities in Maputo and Inhambane provinces of Mozambique (Tessema et al. [2020]). There were 52 samples overall with 26 from each clinic; only samples with data for at least 75 loci were considered for the analysis. From these samples, naïve COI estimates (60% polyclonal samples with maximum COI of 6) and subsequently estimates of population allele frequencies adjusted for COI were calculated. We initially set *M* = 1 and used likelihood-ratio statistics to test a null hypothesis *H*_0_: *r* = 0 at significance level *α* = 0.05 (with the procedure adjusted for a one-sided test). For comparison, Jaccard similarity was used as an IBS metric; Figure 6 displays results from both methods. Dcifer results indicated that the majority of samples were unrelated, and that related samples were mostly from the same clinic. The IBS metric also picked up very highly related pairs, but, apart from those, it was more difficult to distinguish related samples from background. Some samples appeared to be less related to all the other ones in IBS results, and some - more (stripe-like patterns in the lower triangle); these single-sample relatedness levels correlated with estimated COI (e.g. all the lighter “stripes” corresponded to monoclonal samples) highlighting the fact that IBS similarity is strongly influenced by COI, which obscures contribution of descent. For related samples, we also estimated *M′* (the number of related strain pairs) and *r_total_* (overall relatedness). For these samples, 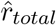 ranged between 0.105 and 1.97 and there were four pairs of samples (all in Maputo), for which 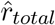 exceeded 1, with estimated COI of 2 in all samples in these pairs and 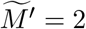 for all such pairs.

**Figure 6:**
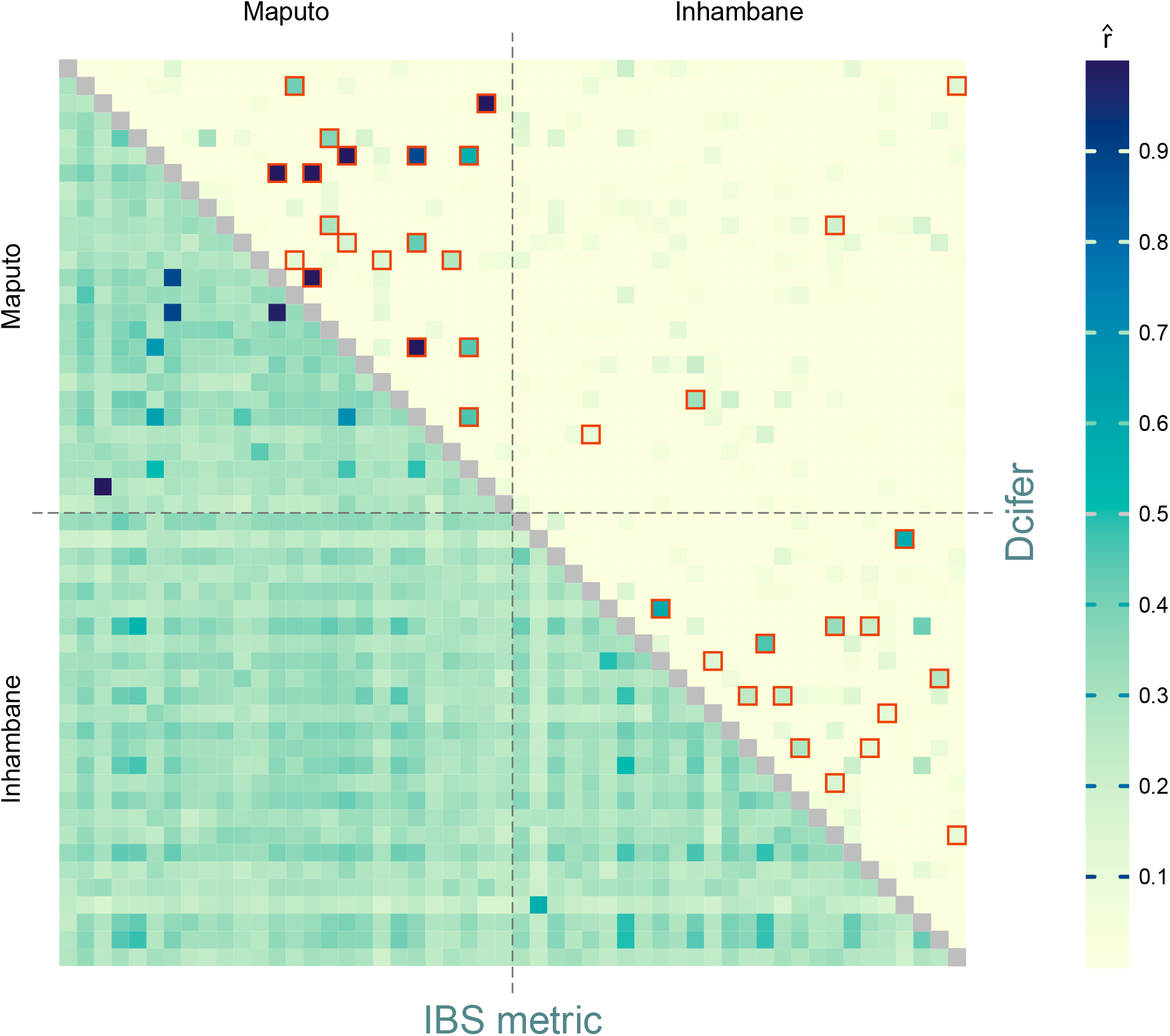
Relatedness between samples from two provinces in Mozambique (Inhambane and Maputo). Lower triangular matrix displays IBS metric results and upper triangular matrix - Dcifer estimates. The color of each matrix entry represents an estimate for two corresponding samples; pairs for which *H*_0_ has been rejected using the Dcifer likelihood ratio test are outlined in red.

We also reevaluated microsatellite data from a previously published dataset, which contained 2585 samples from 29 clinics in four districts in Namibia, using Dcifer to estimate relatedness (Tessema et al. [2019]. These data had 26 loci, and around half of the samples were polyclonal. We assessed the number of related samples within and between clinics (*α* = 0.05), then performed a permutation test to determine which clinic combinations had more related samples than expected by chance. Most of the within-clinic entries had significantly large numbers of related samples (Figure 7). In addition, clinics with geographical proximity had significantly more related between-clinic infections, as illustrated by clusters of darker circles along the diagonal. Rundu DH is a large referral hospital, which could explain relative genetic closeness between samples from this and more geographically distant clinics. Rundu, Nyanganna, and Andara districts are adjacent to each other and Zambezi district is distant from them, which is reflected in the relative lack of relatedness between Zambezi and other districts.

**Figure 7:**
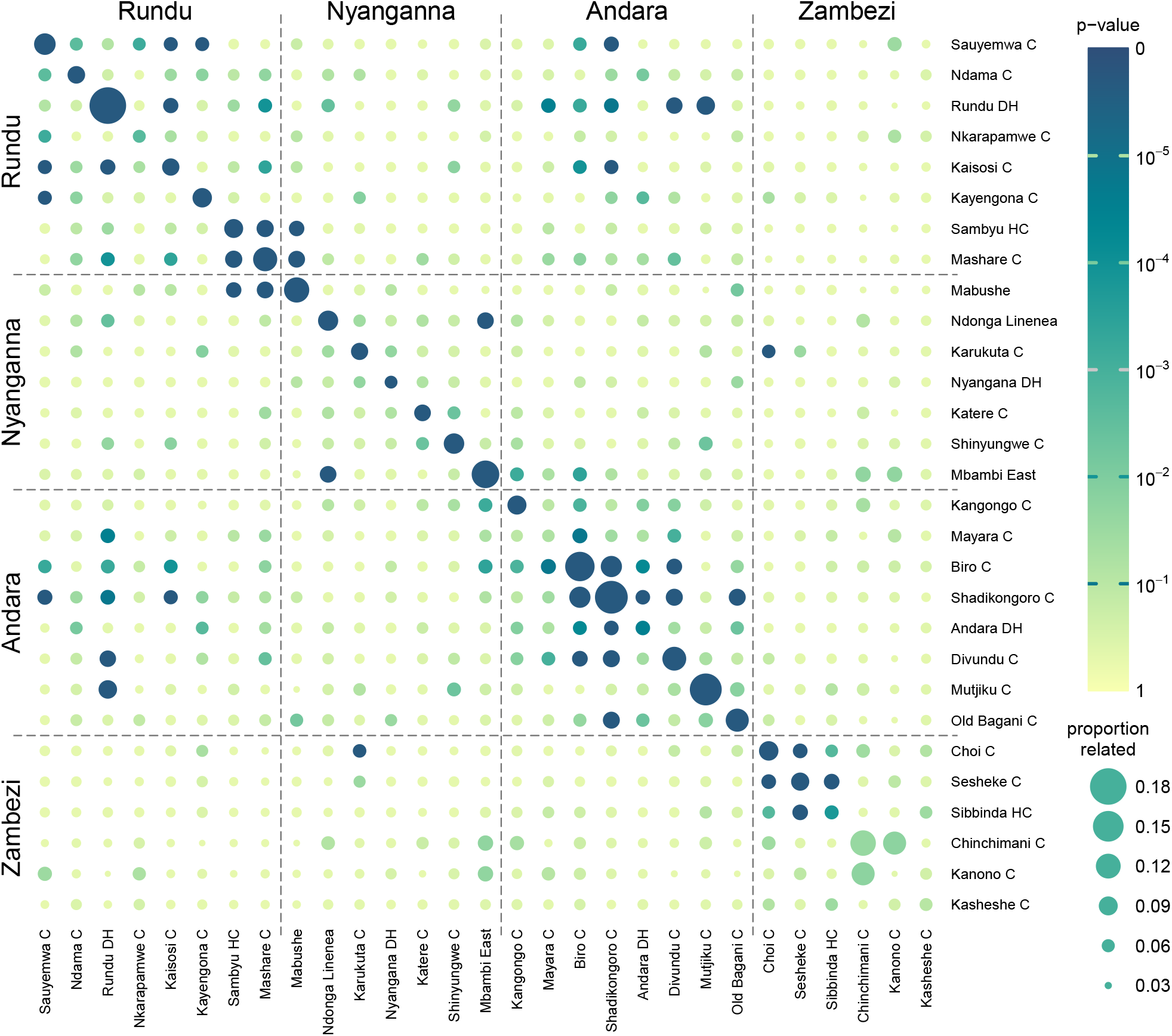
Namibia clinic-level relatedness and permutation test results. Each circle represents a single clinic (on a diagonal) or a two-clinic combination (off-diagonal), with clinics ordered geographically and divided into districts. The color of the circle corresponds to the permutation test’s p-value, and the diameter - to the proportion of related samples within or between the clinics. Permutation distributions for clinics with smaller numbers of samples have larger variance leading to larger p-values - e.g. Chinchimani and Kanono with only 9 samples each have relatively high proportions of related infections but their p-values are not small.

## 4 Discussion

The ability to infer genetic distance between infections is a critical step in translating pathogen genetic data into insight regarding transmission. Despite this, there is a lack of established methods available to infer genetic distance between malaria infections containing multiple parasites, which are the majority in many endemic areas. Options are even more limited when using multiallelic loci, which offer more resolution than biallelic SNPs. The lack of any formal approach has left the community with only ad hoc calculations such as identity by state (IBS), which yield ambiguous results and require extensive efforts to guide any attempt at meaningful inference. In contrast, Dcifer provides relatedness estimates that are based on identity by descent (IBD), interpretable quantities with consistent meaning across studies - regardless of genotyping methods used - with clear implications for ancestry. Importantly, we also show that Dcifer’s statistical power to detect related infections consistently surpasses that of IBS. The method produces reliable measures of uncertainty, and inference obtained from Dcifer versus IBS is more robust to misspecifications of estimated quantities such as COI or population allele frequencies. The R software package implementing Dcifer provides a fast, convenient, and flexible tool that can be easily incorporated into the analysis stream of a wide range of genotyping data to understand transmission.

While Dcifer is designed to work with many types of genotyping data, e.g. biallelic SNPs, multiallelic loci such as microsatellites and microhaplotypes, and any combination thereof, we show that including multiallelic loci results in substantial gains in power to detect related infections. These benefits become more dramatic as COI increases. The result makes intuitive sense, as multiallelic panels provide more within-host strain differentiation and consequently can allow information pertaining to descent to be more easily detected, benefiting relatedness estimation. For example, where two infections with high COI may look similar with biallelic genotyping panels regardless of their level of relatedness (both alleles present at most loci with some diversity), having multiallelic data provides the opportunity to compare these infections more meaningfully. Fortunately, the greater availability of methods to obtain multiallelic data from across the genome make it feasible to generate these data efficiently and in a high throughput manner (LaVerriere et al. [2021], Tessema et al. [2020], Aydemir et al. [2018]).

The concept of relatedness for individual parasites does not extend trivially to polyclonal infections, where strains within and between infections can be related. Dcifer offers an approach that focuses on relatedness between infections as this information is very relevant to transmission. Simulations with imposed intrahost relatedness indicate that the working model achieves its stated goal of capturing interhost relatedness by implicitly downweighting the independent contribution of related strains within a host to comparisons between hosts. Another conceptual issue concerns population allele frequencies, which can affect Dcifer estimates. The foremost question is what constitutes the relevant source population in regards to relatedness between samples and consequently from which data the frequencies should be estimated. If two infections are from communities with different within-community allele frequencies, what are the implications for descent? Dcifer currently assumes the same allele frequencies for both samples but further exploration might be warranted depending on the question of interest. Questions concerning population and scope of the analysis might also arise in regards to potential multiple testing procedures when many pairwise relatedness hypotheses are tested simultaneously. The fact that these hypotheses are not independent should be taken into account when such procedures are considered.

The Dcifer model does not account for linkage disequilibrium and assumes independence of loci. As the malaria genome has relatively short linkage disequilibrium segments, loci independence can be assumed up to a reasonably large number of loci for a correspondingly designed genotyping panel. If, however, the panel has loci which are likely to be linked, e.g. those selected to be in close proximity or for a large number of loci, the independence assumption would no longer hold, which could result in anti-conservative inference. Another limitation is that the model currently does not account for genotyping errors. Future modification could explicitly incorporate the error process via an appropriate model or assess how a specific error process affects the estimates and inference beyond the explorations we have performed here. Another potential venue for further work is developing an MLE estimator for *r_total_* directly as this might become a commonly used summary. A direct estimator might be more efficient, would have the properties of MLE, and would require less processing time. Other future directions could explore alternative inferential approaches, including a non-parametric bootstrap, where loci data would be sampled with replacement. In that case, the fact that variables associated with different loci are not identically distributed, and therefore loci might not be equally informative, would need to be addressed.

With potential to facilitate understanding of relatedness structure from unphased genetic data, including multiallelic loci, Dcifer can provide a vital link in the analytical process leading to better understanding of malaria transmission dynamics. While we have demonstrated the utility of this method for *Plasmodium* infections here, Dcifer may be useful in analyses of other organisms that undergo sexual recombination and where polyclonal infections are encountered, such as shistosomiasis, filarial disease, and soil transmitted helminths (Brouwer et al. [2001], Churcher et al. [2008]). With the ability to incorporate most types of genetic data, rapid computation, and readily available inference, Dcifer may prove to be an important tool in the analytical toolbox for obtaining epidemiologic insight from pathogen genetics.

## Data availability

File Supplemental Data 1 contains microhaplotype data from Mozambique. Microsatellite data from Namibia are publicly available at https://elifesciences.org/articles/43510/figures#supp1.

## Acknowledgments

We thank Nicholas Hathaway, Sofonias Tessema, Francisco Saute, and Pedro Aide for generating and sharing microhaplotype data from South Mozambique and for their assistance with extracting relevant information. We also thank the authors of Tessema et al. [2019] for making data from Namibia publicly available. We are grateful to Aimee Taylor for fruitful discussions regarding relatedness and genetic distance and inspiring this work with very clearly presented concepts in Taylor et al. [2019].

## Supplementary Materials

### S.1 A note on model assumptions

The assumption of independence of all interhost *IBD* variables at a given locus, while convenient, does not at first appear fully aligned with the goal of capturing interhost relatedness, as it leaves out a potential source of relatedness that could be present in the dependent case. For example, *r*_*x*_1_,*y*_1__ and *r*_*x*_2_,*y*_1__ could be both positive and still satisfy a “no intrahost relatedness” assumption (*r*_*x*_1_,*x*_2__ = 0) with a certain type of dependence between *IBD*_*x*_1_,*y*_1_,*t*_ and *IBD*_*x*_2_,*y*_1_,*t*_. For a simplest special case of such dependence, consider the first two strains in the first sample being the unrelated parents of the first strain in the second sample; then

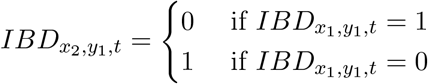

The general case can be described as multiple strains in one infection (say, strains *i* = 1, …, *m*) being unrelated to each other while all related to a strain in another infection (strain *j*). In that case, the sum 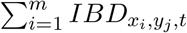 of corresponding *IBD* variables does not exceed 1: 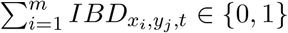, and the working model will still be able to capture 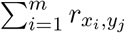 by treating 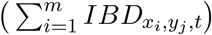 as a single binary variable and thus arriving at the desired estimate of overall relatedness or individual relatedness estimates within the framework.

### S.2 Constructing the likelihood

We start with the simplest case of *M* = 1, ***r*** = *r*. Then

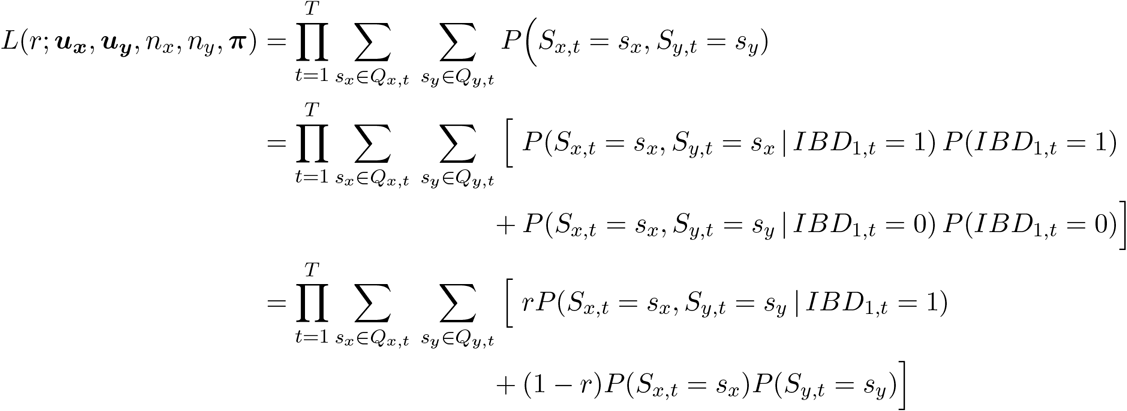

If *IBD*_1,*t*_ = 0, *S_x,t_* and *S_y,t_* are independent, and any matching of the alleles between them occurred by chance. Using multinomial distributions, *P*(*S_x,t_* = *s_x_*) *P*(*S_y,t_* = *s_y_*) = *g*(*s_x_*; *n_x_*, (*π_t_*)) *g*(*s_y_*; *n_y_*, (*π_t_*)). If *IBD*_1,*t*_ = 1, one of the shared alleles *a_t,i_* ∈ *u_xy,t_*, where *u_xy,t_* = *u_x,t_* ⋂ *u_y,t_*, is matching because of shared ancestry (if there are no shared alleles, *IBD*_1,*t*_ cannot be 1). Fixing this allele in place and allowing other alleles in *s_x_* and *s_y_* to “fill” the rest of the haplotypes, we get

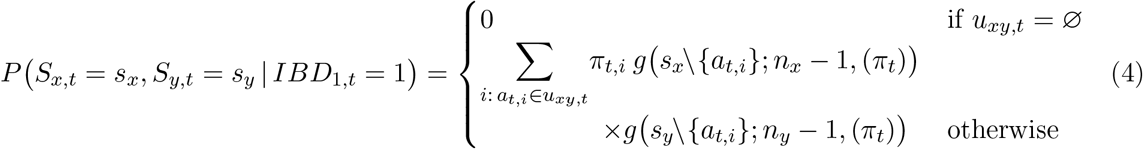

When 1 < *M* ⩽ |*s_xy,t_*| and 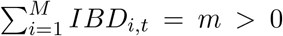, the “fixed” portion of the interhost strain pairs can itself be thought of in terms of a multinomial distribution *Multinom*(*m*, *π*_*t*,1_, …, *π_t,K_t__*), extending the base *M* = 1 case. We illustrate the process with *M* = 2:

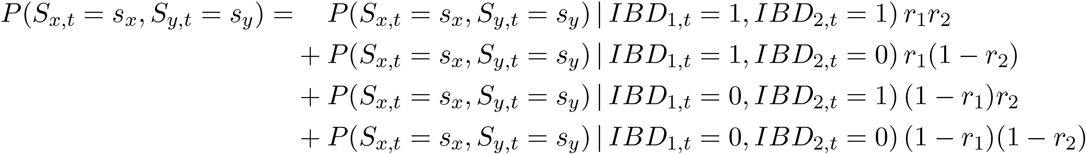

*P*(*S_x,t_* = *s_x_*, *S_y,t_* = *s_y_*) | *IBD*_1,*t*_ + *IBD*_2,*t*_ = 1) is equal to (4) and

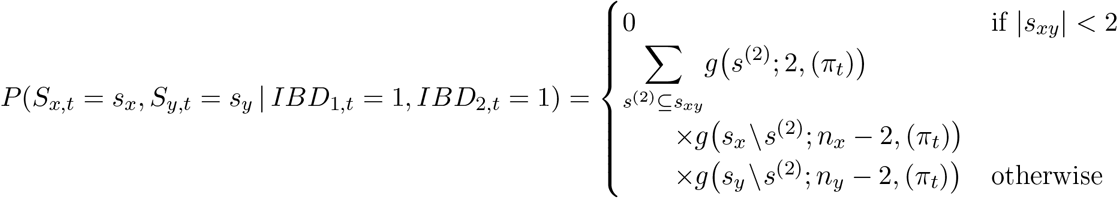

Conditional probability *P*(*S_x,t_*, *S_y,t_* | *IBD*_1,*t*_, …, *IBD_M,t_*) is the same for all 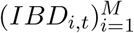 sequences with the same 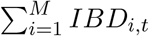, which makes calculations considerably faster.

### S.3 Breaking down the support curve

To get some intuition on the behavior of the support curve and how it is affected by the sample data, COI, and population allele frequencies, we explore the likelihood for a single locus (sample size is 1) when *M* = 1 (***r*** = *r*). For locus *t*,

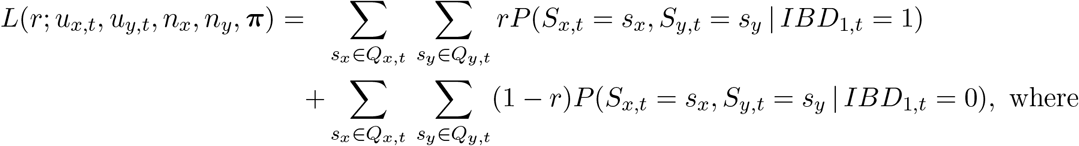

*P*(*S_x,t_* = *s_x_*, *S_y,t_* = *s_y_* | *IBD*_1,*t*_ = 1) is as in (4) and

*P*(*S_x,t_* = *s_x_*, *S_y,t_* = *s_y_* | *IBD*_1,*t*_ = 0) = *g*(*s_x_*; *n_x_*, (*π_t_*)) *g*(*s_y_*; *n_y_*, (*π_t_*)). Let

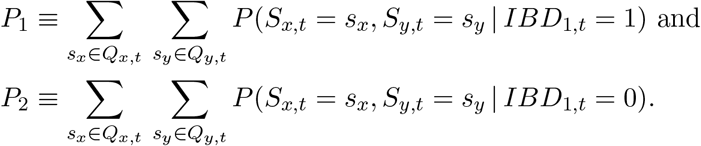

Then the log-likelihood for locus *t* can be written as

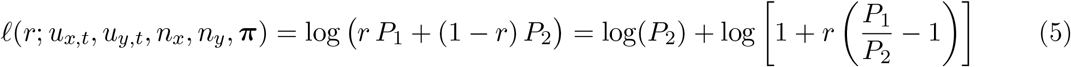

Since logarithmic function is monotonic, the log-likelihood for a single locus is a monotonic function of *r* and a maximum likelihood estimate is either 0 or 1. Thus we can explore when the function is increasing or decreasing and what is the shape of the support curve (note that the function is concave as the second derivative is negative or zero). First, consider the case when *P*_1_/*P*_2_ is close to 1. Then (*P*_1_/*P*_2_ – 1) is small and log [1 + *r* (*P*_1_/*P*_2_ – 1)] ≈ *r* (*P*_1_/*P*_2_ – 1) for any *r*, which means that the log-likelihood is approximately linear.

To compare *P*_1_ and *P*_2_, we can look at their components - allele frequencies and multinomial coefficients. A simple case with a single shared allele can illustrate their comparison but the concept readily extends to multiple shared alleles. *P*_2_ has an extra factor of *π_i_*, where *a_i_* is a shared allele; on the other hand, it has higher factorials - depending on a combination of these factors, *P*_1_ is either less than *P*_2_ (log-likelihood is decreasing) - or greater (log-likelihood increasing). Smaller *π_i_* contributes to a greater *P*_1_/*P*_2_ ratio (an argument for relatedness), as does a greater number of shared alleles (|*u_xy,t_*|). This can also elucidate the “non-linear” cases where *P*_1_/*P*_2_ is not close to 1: if the frequencies of shared alleles are very small, the ratio is high (strong evidence of relatedness), and if there are no shared alleles, *P*_1_ = 0 and log-likelihood goes to –∞ at *r* = 1. This stronger evidence in support or against independence in a locus increases contribution of that locus to the overall likelihood thus having a greater effect on the relatedness estimate. Furthermore, its effect on likelihood-ratio-based inference can be even greater; for example, a shared allele with a very small population frequency in a single locus can lead to rejecting *H*_0_: *r* = 0 even when the estimate itself is low. For practical implications, this could underline the importance of allele frequencies estimation: e.g. estimates suffering from biases in data selection, such as data not being representative of population in terms of allele frequencies, can significantly affect downstream results.

### S.4 Limitations of bootstrap-based asymmetric confidence intervals

This section contains notes on constructing confidence intervals using bootstrap and applications of this method to Dcifer although it is not implemented in Dcifer directly for the reasons described below. Let 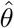 be an estimate of a parameter *θ*, for which we want to find a 1 – *α* confidence interval (CI). Let *CI_lo_* and *CI_up_* be the values at the endpoints of such *CI*:

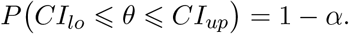

Fact:

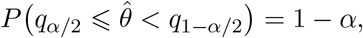

where *q*_*α*/2_ and *q*_1–*α*/2_ are *α*/2’th and (1 – *α*/2)’th quantiles of the sampling distribution. Then

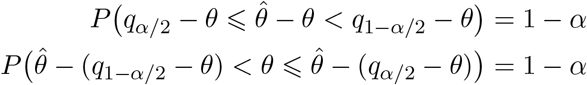

and

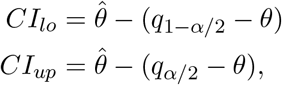

where *q*_*α*/2_, *q*_1–*α*/2_, and *θ* are unknown.

Suppose we use bootstrap to approximate *q*_*α*/2_ – *θ* with 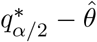 and *q*_1–*α*/2_ – *θ* with 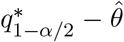, where 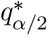 and 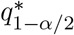 are *α*/2’th and (1 – *α*/2)’th quantiles of the bootstrap distribution. Then

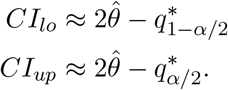

Inverting the quantiles for the CI makes intuitive sense: if a sampling distribution is skewed to the right, and 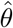 tends to overestimate *θ*: *θ* – *q*_*α*/2_ < *q*_1–*α*/2_ – *θ*, then *CI_lo_* should be further from 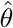 than *CI_up_*. Validity of such inference rests on the bootstrap principal assumption that the distribution of 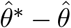 is similar enough to the distribution of 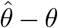. However, in our case with bounded support (0 ⩽ *θ* ⩽ 1), these distributions can be quite different. That principal assumption would hold in the midrange, provided that the sample size is large enough, when bootstrap-based CI’s are reasonably symmetric; in that case, however, they would be very similar to Wald and likelihood-ratio-based intervals.

### S.5 Population allele frequencies and observed allele counts

Let *N* be the number of samples (infections) with *n*_1_, …, *n_N_* parasite strains in them (COI). For a given locus *t* and a given allele *a_t,k_* with population frequency *π_t,k_*, let *b* = (*b*_1_, …, *b_N_*), *b_i_* ∈ {0, 1) be a binary sequence of indicators of whether that allele is present in each sample. Then, for a sample *i*,

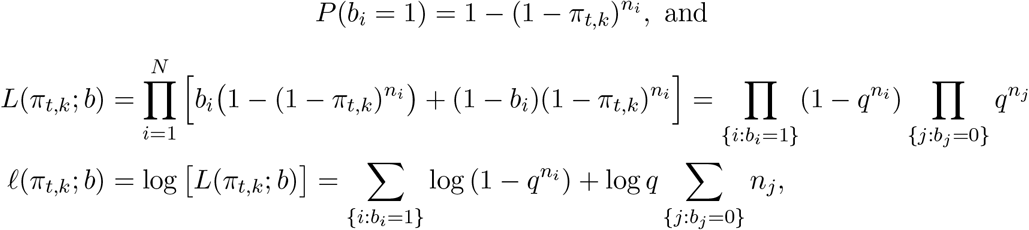

where *q* = 1 – *π_t,k_*.

If *n*_1_ = … = *n_N_* = 1 (all the infections are monoclonal), 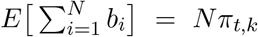; otherwise 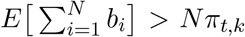. Naive estimation of allele frequencies as normalized proportions of number of samples with each allele at a given locus will result in underestimating frequencies of common alleles and overestimating rare allele frequencies. In turn, that would lead to higher heterozygosity, and, consequently, to overestimating relatedness between infections. The reason for such overestimation is that sharing of common alleles would be considered less likely to have occured by chance (as opposed to descent) than it actually is. Therefore it is advisable to adjust allele frequencies estimation to account for complex infections.

### S.6 Supplementary figures

**Figure S.1:**
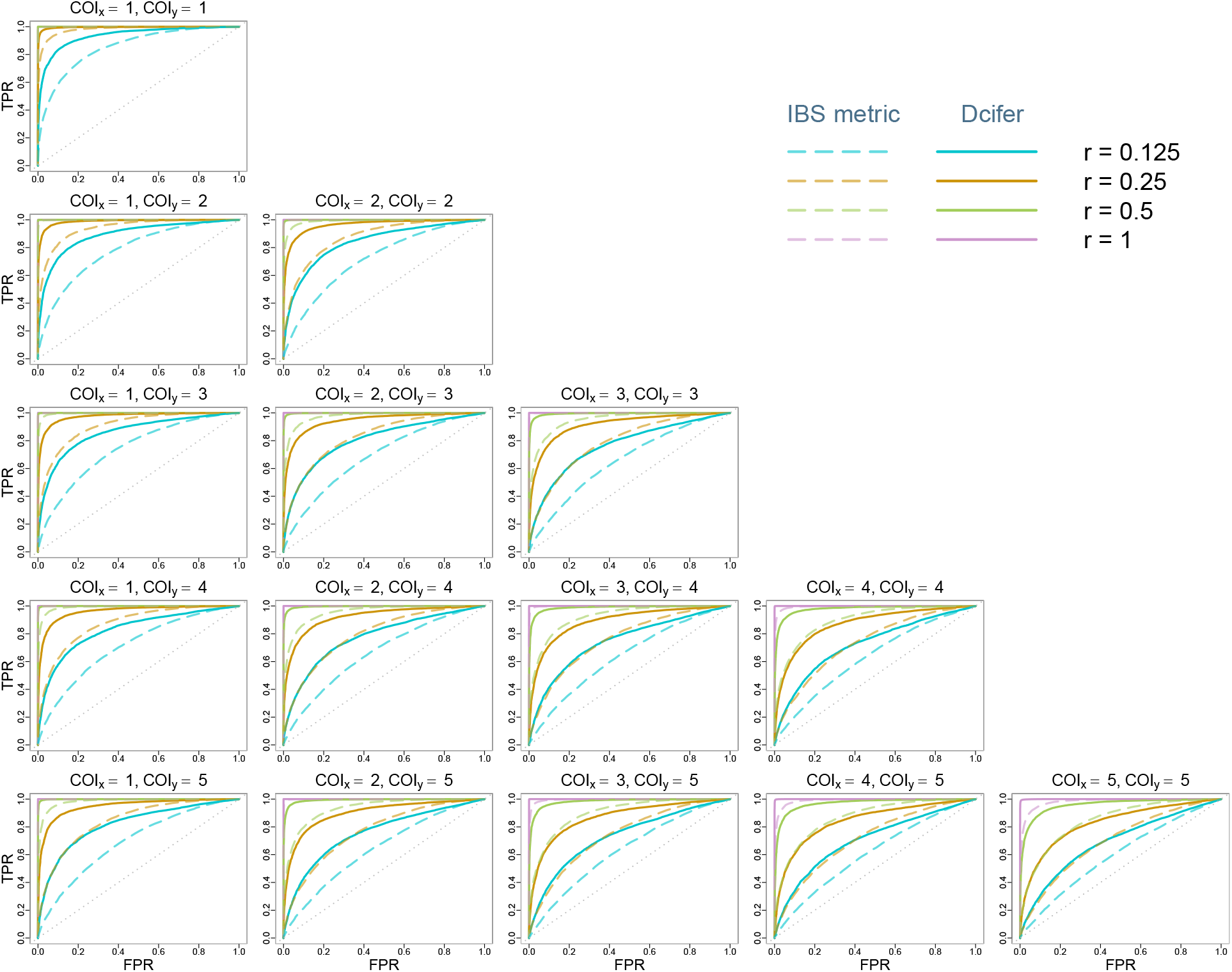
Receiver operating characteristic curves for Dcifer relatedness estimator 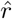 and IBS similarity metric, classification accuracy for related when compared to *r* = 0, using the data presented in Figure 2.

**Figure S.2:**
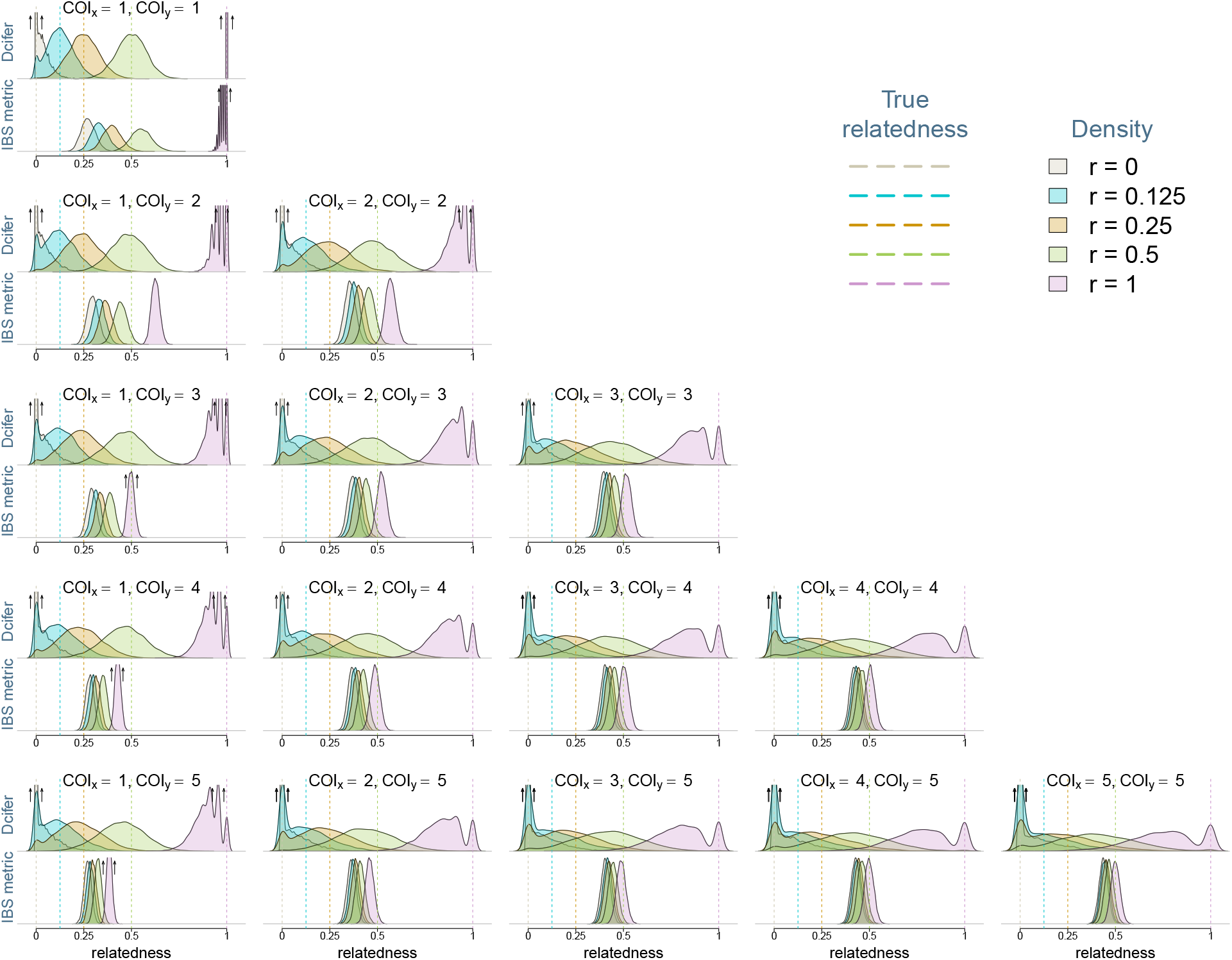
Densities of Dcifer relatedness estimator 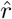 and IBS similarity metric results obtained from data simulated with genotyping errors (*ϵ* = 0.05, λ = 0.01) using a panel of 91 microhaplotypes. Simulations were performed for five values of *r* and for COI combinations ranging between 1 and 5. Estimated COI and population allele frequencies were used for Dcifer.

**Figure S.3:**
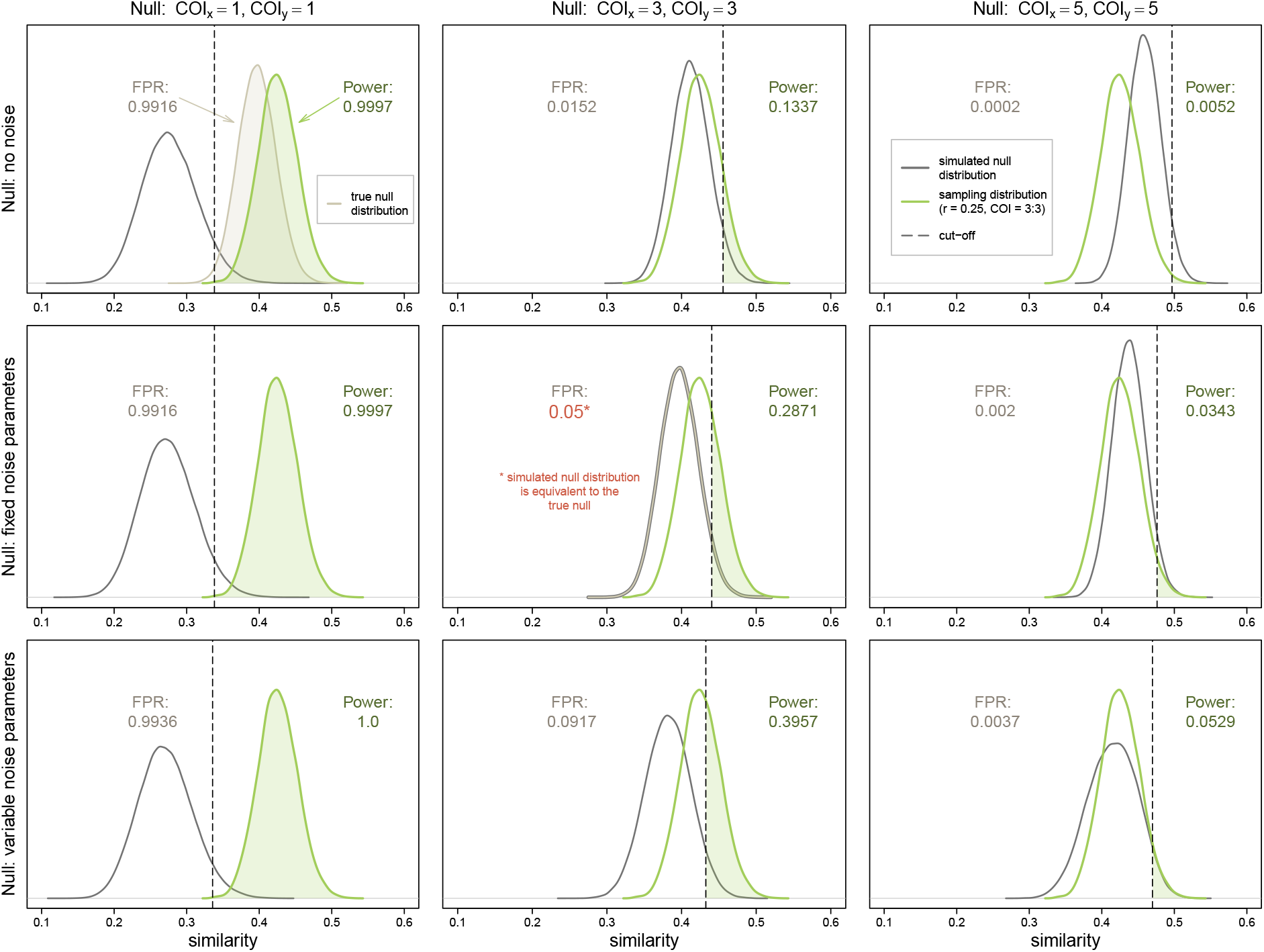
Hypothesis testing with different reference (simulated null) distributions for the IBS metric applied to related infections (*r* = 0.25) with COI of 3. Vertical dotted line in each panel represents a rejection cut-off determined by the 0.95-quantile of the corresponding null (the area under the density curve in dark grey to the left of the line is equal to 0.95). The green curve of the sampling distribution and the light beige curve of the true null displayed in the top left panel are the same for each panel. Note the narrow range of the values (x-axis range represents only part of the [0, 1] support). Depending on the assumed null distribution, the power ranges between 0.0052 and 1 and the false positive rate (FPR) - between 0.0002 and 0.9936, highlighting how sensitive this inferential approach is to assumptions about COI and error, in contrast to the likelihood ratio approach available using Dcifer.

**Figure S.4:**
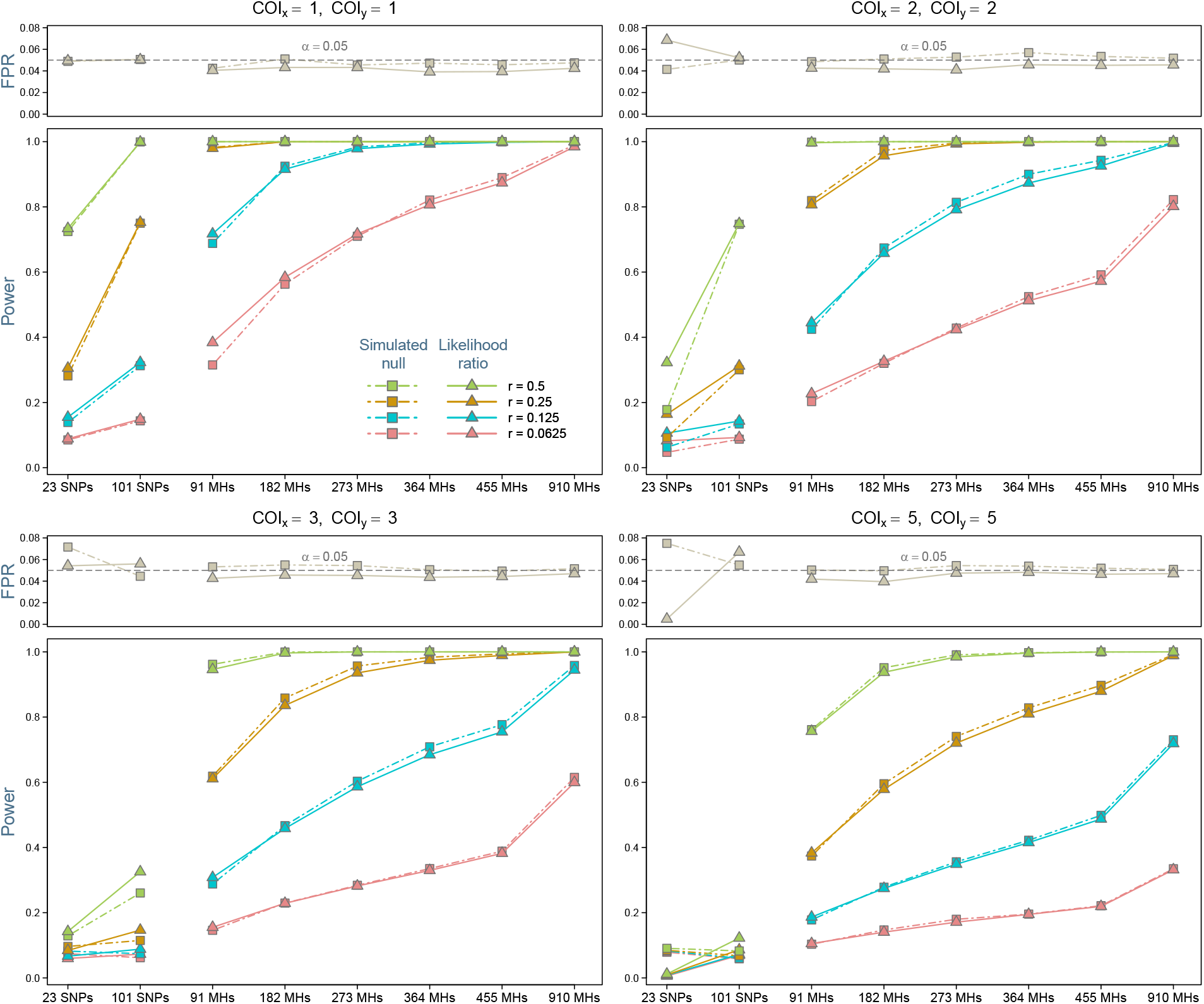
Comparison of two inferential approaches within Dcifer: simulated null distributions vs likelihood ratio. False positive rate and statistical power of a test *H*_0_: *r* = 0 at significance level *α* = 0.05 are shown. Simulations were performed with genotyping error, and COI were estimated from these data.

**Figure S.5:**
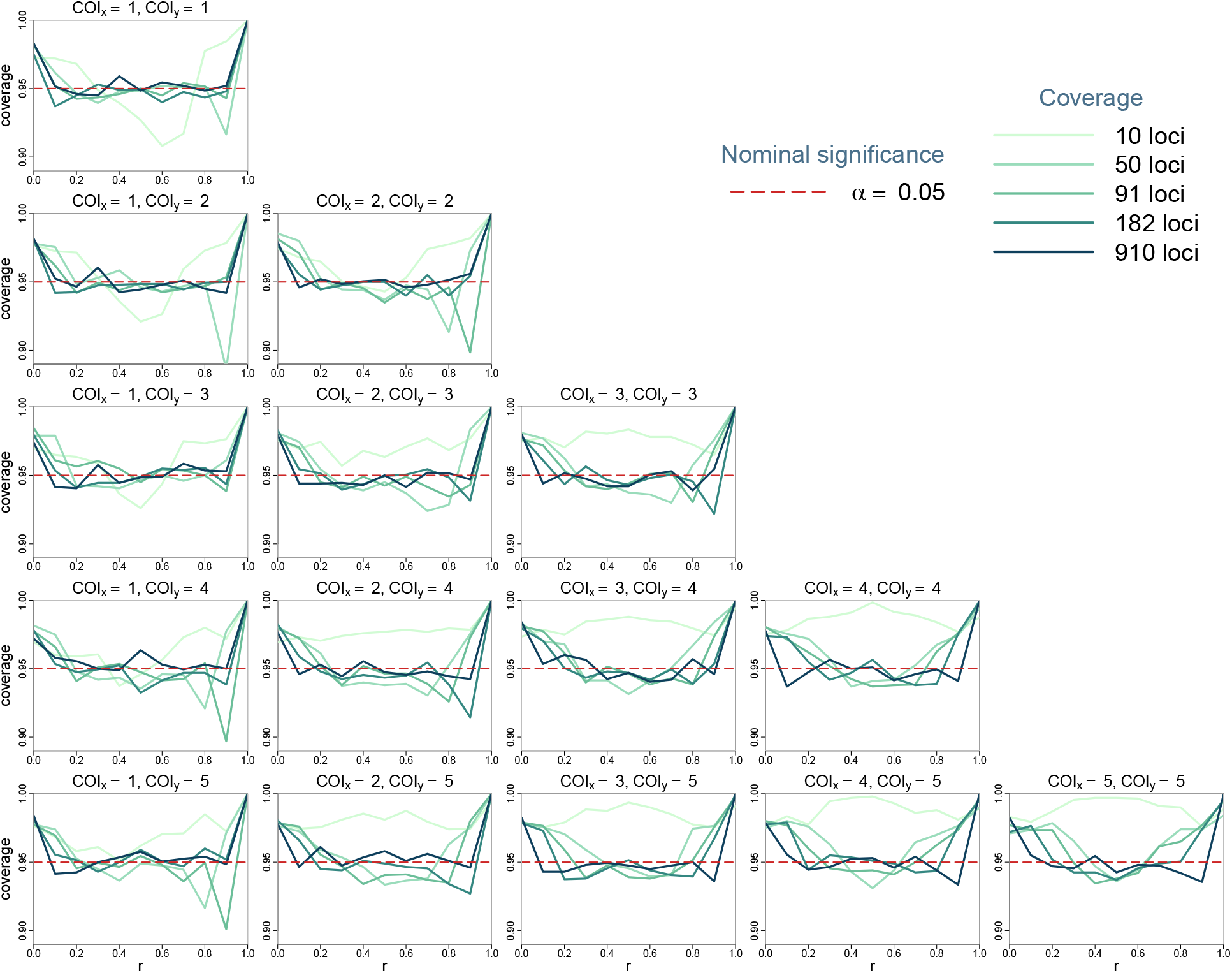
Coverage for likelihood-ratio-based 95% confidence intervals produced by Dcifer.

**Figure S.6:**
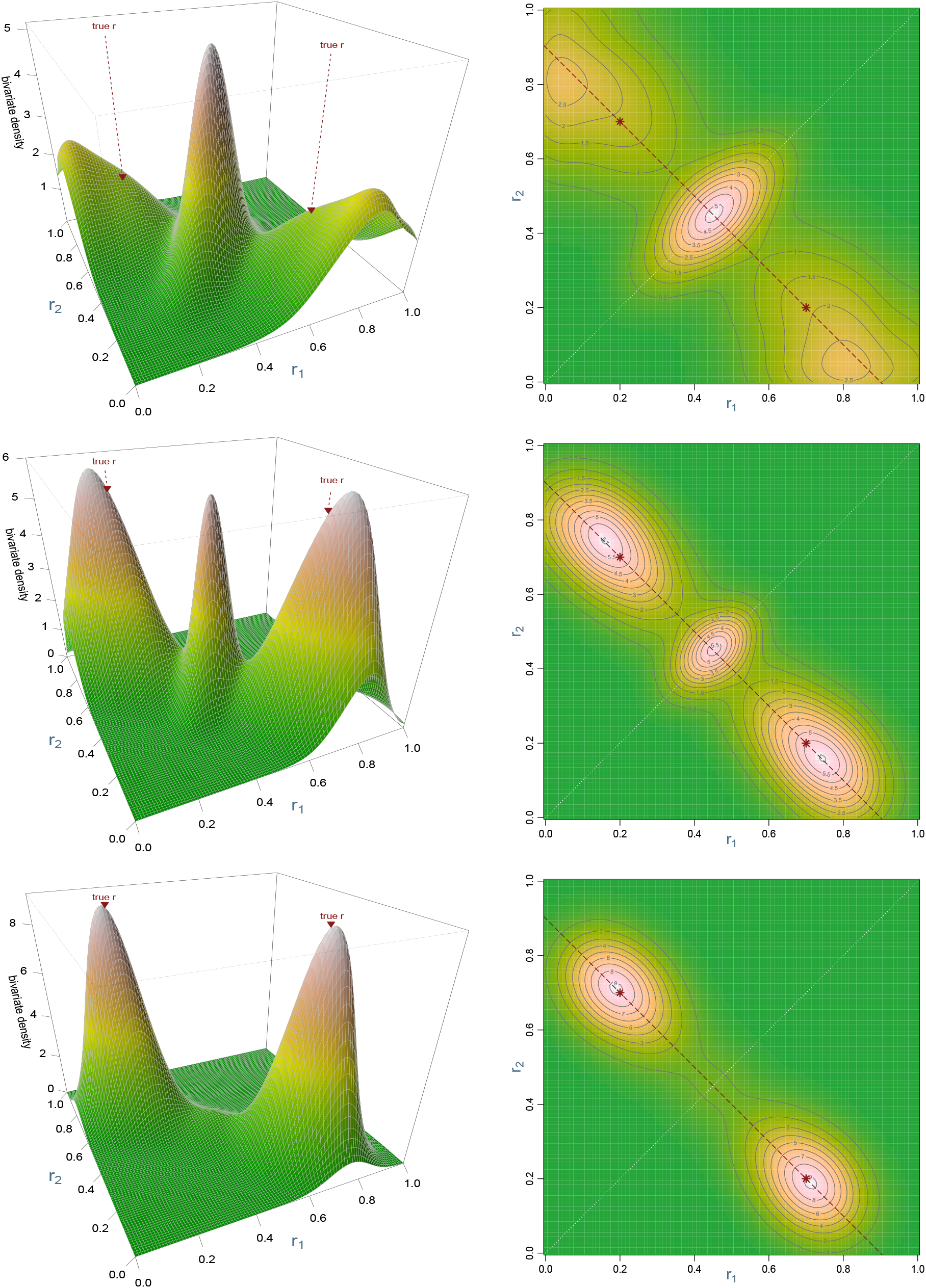
Sampling distribution for *M* = 2, *r*_1_ = 0.2, *r*_2_ = 0.7, *n_x_* = *n_y_* = 3. True values are indicated by red stars on the contour plots; red dotted line represents *r_total_* = *r*_1_ + *r*_2_. Top row: 91 loci, middle row: 273 loci, bottom row: 910 loci. We can see two tendencies in the estimation of multiple parameters: one is for the estimates to be equal and another - to separate them further, pulling some estimates toward 0 or 1.

**Figure S.7:**
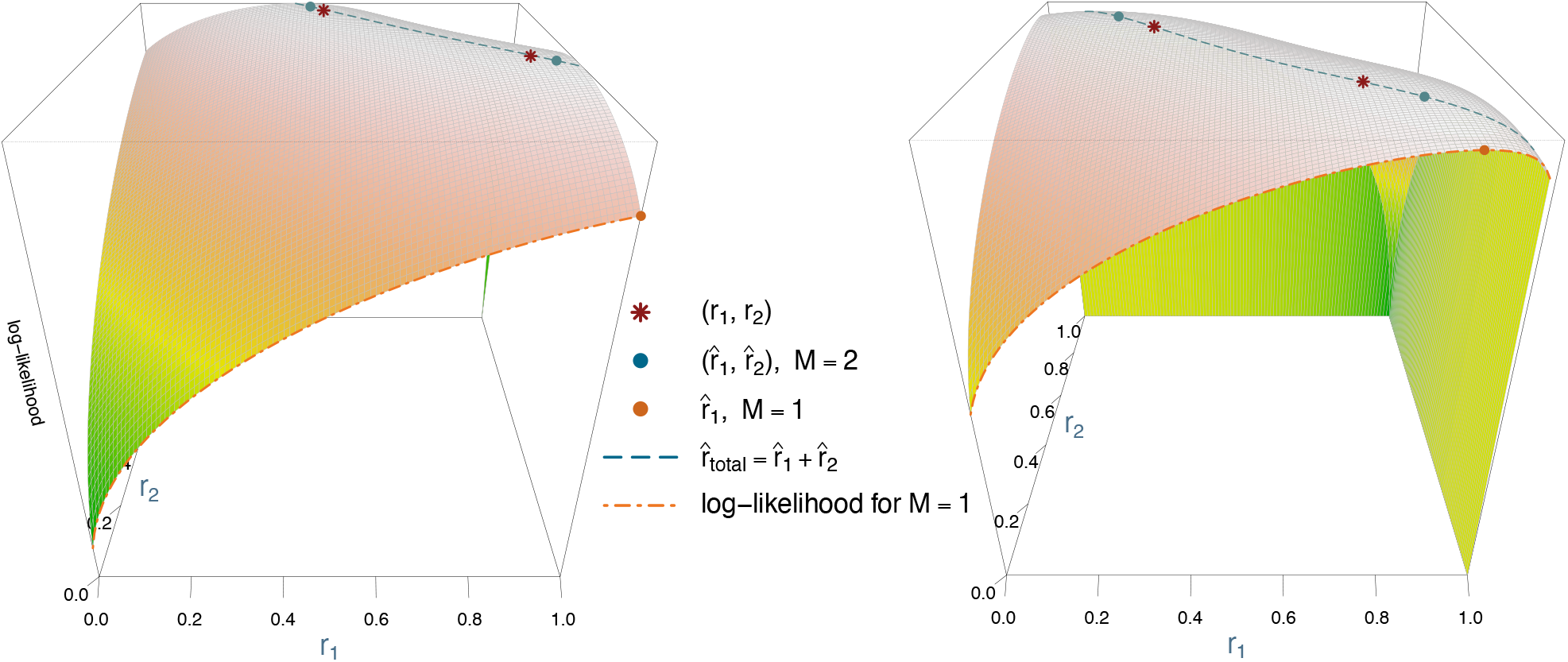
Examples of log-likelihood surfaces for two pairs of related infections with COI of 3, *M* = 2. The true parameter values (*r*_1_, *r*_2_) are (0.48, 0.92) in the left panel and (0.31, 0.74) in the right.

**Figure S.8:**
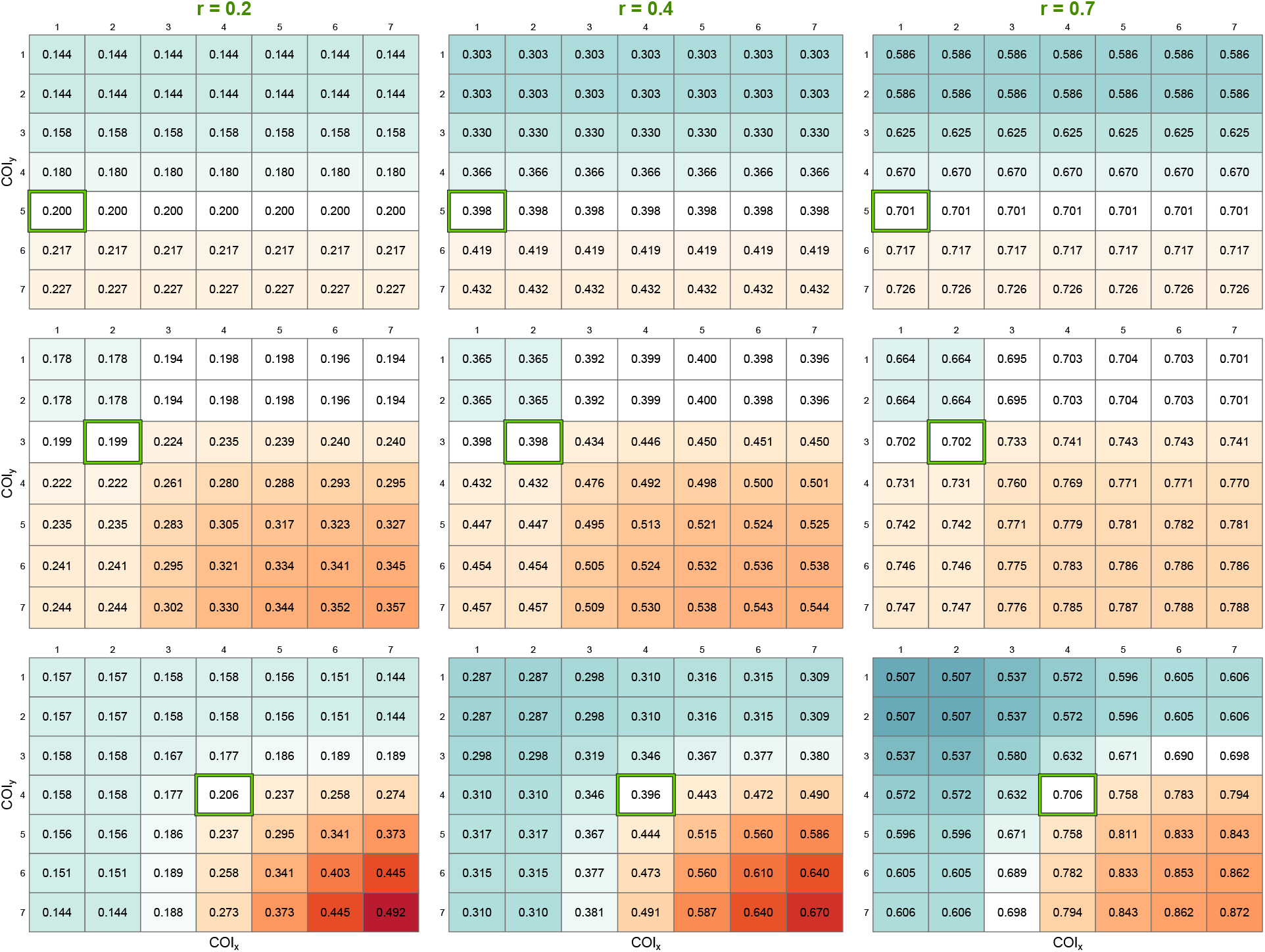
Effect of COI misspecifications on relatedness estimates. Each panel represents a matrix whose entries are average estimates obtained with a corresponding COI combination. True COI is outlined in green; background color of each entry corresponds to the deviation from the true *r* value.

**Figure S.9:**
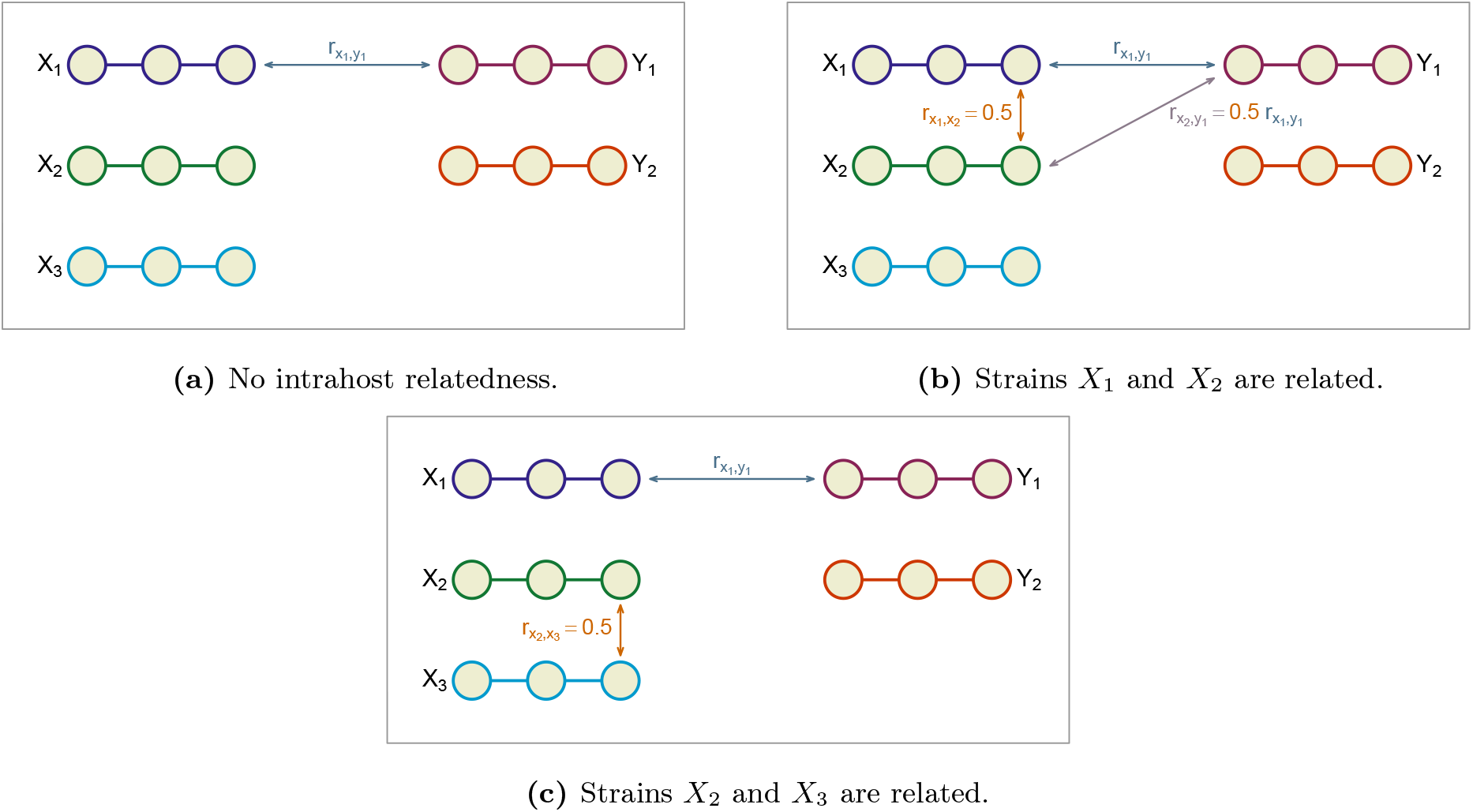
Diagram of intrahost relatedness simulations using an example with COI of 3 and 2. Three scenarios are compared, each with the same level of relatedness between strains represented by *X*_1_ and *Y*_1_, which is not changed by the induced intrahost relatedness in (b) and (c). Horizontal chains of connected circles represent haplotypes.

**Figure S.10:**
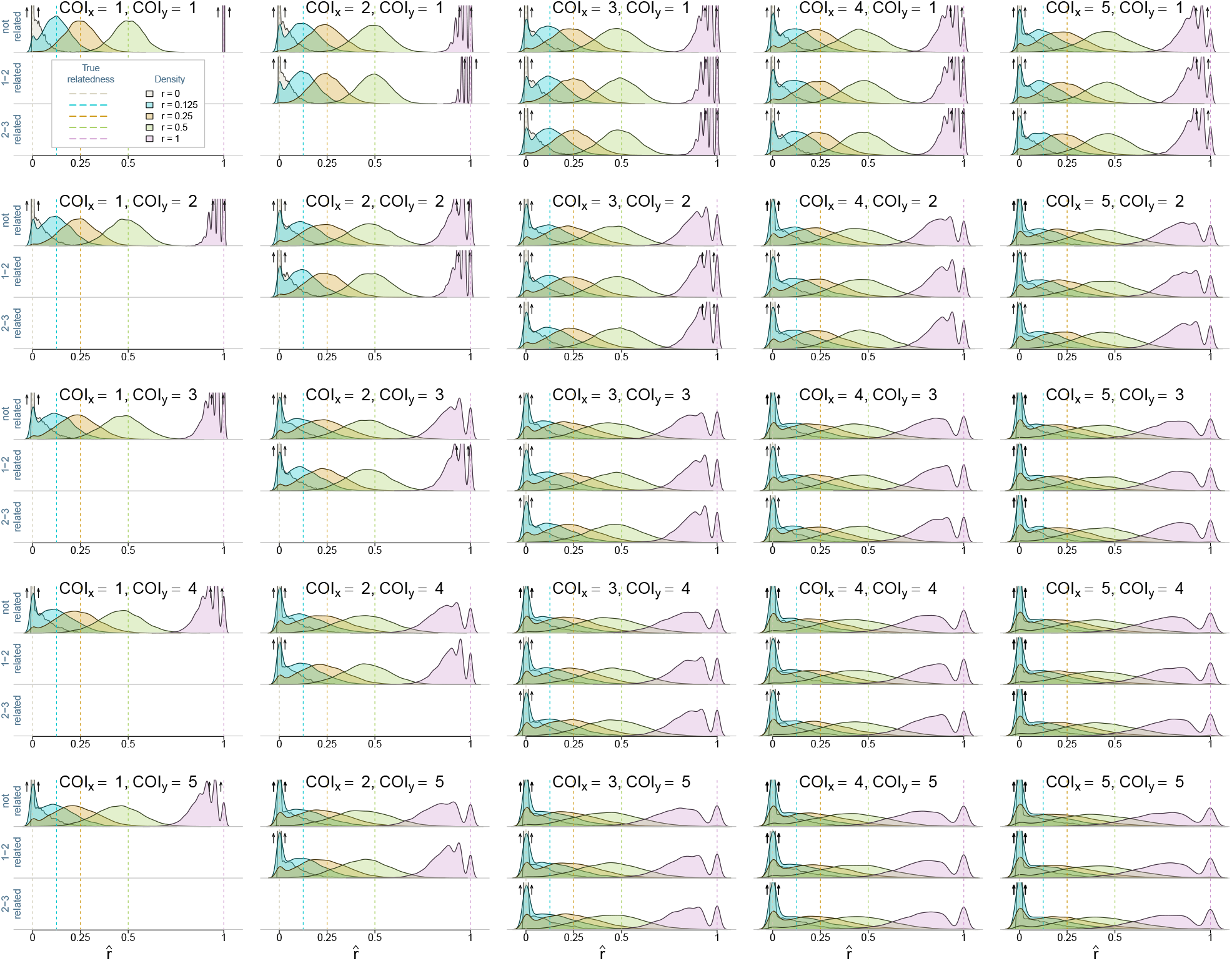
Comparison of sampling distributions across three intrahost relatedness scenarios: 1) no intrahost relatedness, 2) strains *X*_1_ and *X*_2_ are related, 3) strains *X*_2_ and *X*_3_ are related (as illustrated in Figure S.9).

